# Identifying commonalities between cell lines and tumors at the single cell level using Sobolev Alignment of deep generative models

**DOI:** 10.1101/2022.03.08.483431

**Authors:** Soufiane M.C. Mourragui, Joseph C. Siefert, Marcel J.T. Reinders, Marco Loog, Lodewyk F.A. Wessels

## Abstract

Preclinical models are essential to cancer research, however, key biological differences with patient tumors result in reduced translatability to the clinic and high attrition rates in drug development. Variability among and between patients, preclinical models, and individual cells obscures commonalities which could otherwise be exploited therapeutically. To discover the shared biological processes between cell line models and clinical tumors we developed **Sobolev Alignment**, a computational framework which uses deep generative models to capture non-linear processes in single-cell RNA sequencing data and kernel methods to align and interpret these processes. We show that our approach faithfully captures shared processes on a set of three synthetic datasets. Exploiting two large panels of untreated non-small cell lung cancer cell lines and patients, we identify the similarities between cell lines and tumors and show the conservation of key mitotic and immune-related pathways. Employing our approach on a large in-vitro perturbation screen, we show that processes captured by our method faithfully recapitulate the known modes of action of clinically approved drugs and allow investigation into the mode of action of an uncharacterized drug.

## Introduction

Synthetic model systems, like cell lines, offer a highly cost-effective and versatile way to study human biology. In the case of cancer, the facility to study these model systems in a wide array of conditions renders them particularly attractive for drug screening^1–3^. When compared to human tumors, however, these benefits are over-shadowed by intrinsic limitations such as the lack of a vasculature or micro-environment^4–6^. Consequently, understanding the biological differences between pre-clinical models and patients is key to improving the translation of findings from basic research to clinical implementation^7,8^.

Several computational studies have already attempted to characterize the molecular differences between cell lines and patients. The first category of approaches consist of designing machine learning tools to capture the common information relevant for transferring biomarkers of drug response^9–14^. A second category of approaches compares the genomic landscapes of cell lines and tumors in an unsupervised fashion^15–17^. The latter studies have notably highlighted the existence of key differences in molecular profiles and identified clear differences between certain cancer cell lines and their tissues of origin.

These insightful studies are based on bulk RNA-seq data, where gene expression is averaged over thousands of cells. Single cell sequencing technologies, like single-cell RNA sequencing (scRNA-seq), provide a more fine-grained view by measuring gene expression profiles for each individual cell. In this work, we set out to assess the similarities and differences in transcriptional profiles between cell lines and tumors at single cell resolution. Specifically, we focus on cell line cultures^18,19^ and tumors extracted from lung cancer patients^20^, as non-small cell lung cancer (NSCLC) cell lines have been shown to markedly drift from their tissue of origin^16^. Using a panel of lung cancer tumors and a panel of cell lines, we first show that the differences between cell lines and tumors cannot be modeled as a classical batch effect. As previous methods comparing cell lines to tumors were not designed for scRNA-seq data^11,13^, we developed a novel computational approach, **Sobolev Alignment (SA)**, which mobilizes recent advances in large-scale kernel-based machine learning^22–24^ and deep-learning-based probabilistic modelling of scRNA-seq profiles^25–27^. We show that the application of SA to three simulated single cell datasets accurately captures the (constructed) shared biology between datasets. Using SA, we then set out to characterize the shared and distinct biology between NSCLC cell lines and tumors. Amongst others, we showed the conservation of a wide array of immune-related processes. By exploiting the biological processes shared between cell lines and tumors, we demonstrated that our approach recapitulates known modes of action of four clinically approved drugs, Finally, we analyzed the perturbation triggered by a drug of unknown mechanism.

## Results

### Divergence between gene expression profiles of Non-Small Cell Lung Cancer (NSCLC) cell lines and tumors obscures cell-type definition

Cancer cell lines established from biopsies of NSCLC patients are usually assumed to correspond to homogenous cell populations classified as differentiated epithelial cells. Following this assumption, integrating single-cell profiles measured on NSCLC cell lines and tumors should yield a perfect co-clustering^28–32^. This assumption is however challenged by recent studies which highlight significant differences in gene expression patterns^11,16^, mostly resulting from the lack of micro-environment in cell lines (**Figure 1**A). To assess this cell-type drift and evaluate the ability of existing batch-effect correction tools to correct it, we employed three state-of-the-art approaches^33,34^: Seurat v3^29^, Harmony^35^ and LIGER^32^ (Methods). For tumors, we use a panel of 208,506 cells from 58 NSCLC cancer patients at different disease stages^20^, including 36,467 epithelial cells (**Extended Figure 1**A-B), referred to as the *Kim dataset*. For cell lines, we use a panel of 53,512 single cells from 198 cell lines^18^, including 11,105 single cells from 32 NSCLC cell lines (**Extended Figure 1**C), referred to as the *Kinker dataset*. We characterized cells along two axes: 1) derived from cell lines or tumors, and 2) whether the cells are epithelial NSCLC cells, micro-environment-related cells (tumors), or from a different cancer-type (cell lines). We visualized the degree of co-clustering using UMAP^36^. For both Seurat (**Figure 1**B) and Harmony (**Figure 1**C), we observe no co-clustering of epithelial tumor cells and NSCLC cell line cells. LIGER offered a slightly better co-clustering (**Figure 1**D), with one cluster mixing cell-lines and tumors from the same cell type, although not perfectly. For all three methods, however, most NSCLC cancer cell lines are projected away from the epithelial tumor cells, indicating either a clear difference in gene expression profiles or an inability of current methods to correct this kind of batch effect. When looking at the tumor clustering, we furthermore observe a lack of mixing in epithelial tumor cells, contrasting with a good mixing in tumor micro-environment cells (**Extended Figure 1**D-F). This suggests a profound difference between transcriptional profiles of cell lines and tumor cells, hampering standard single cell batch alignment pipelines severely. To reconcile these differences, we set out to systematically identify and compare biological processes present in cell lines and tumors and use them to align the two datasets (**Figure 1**E).

**Figure 1.**
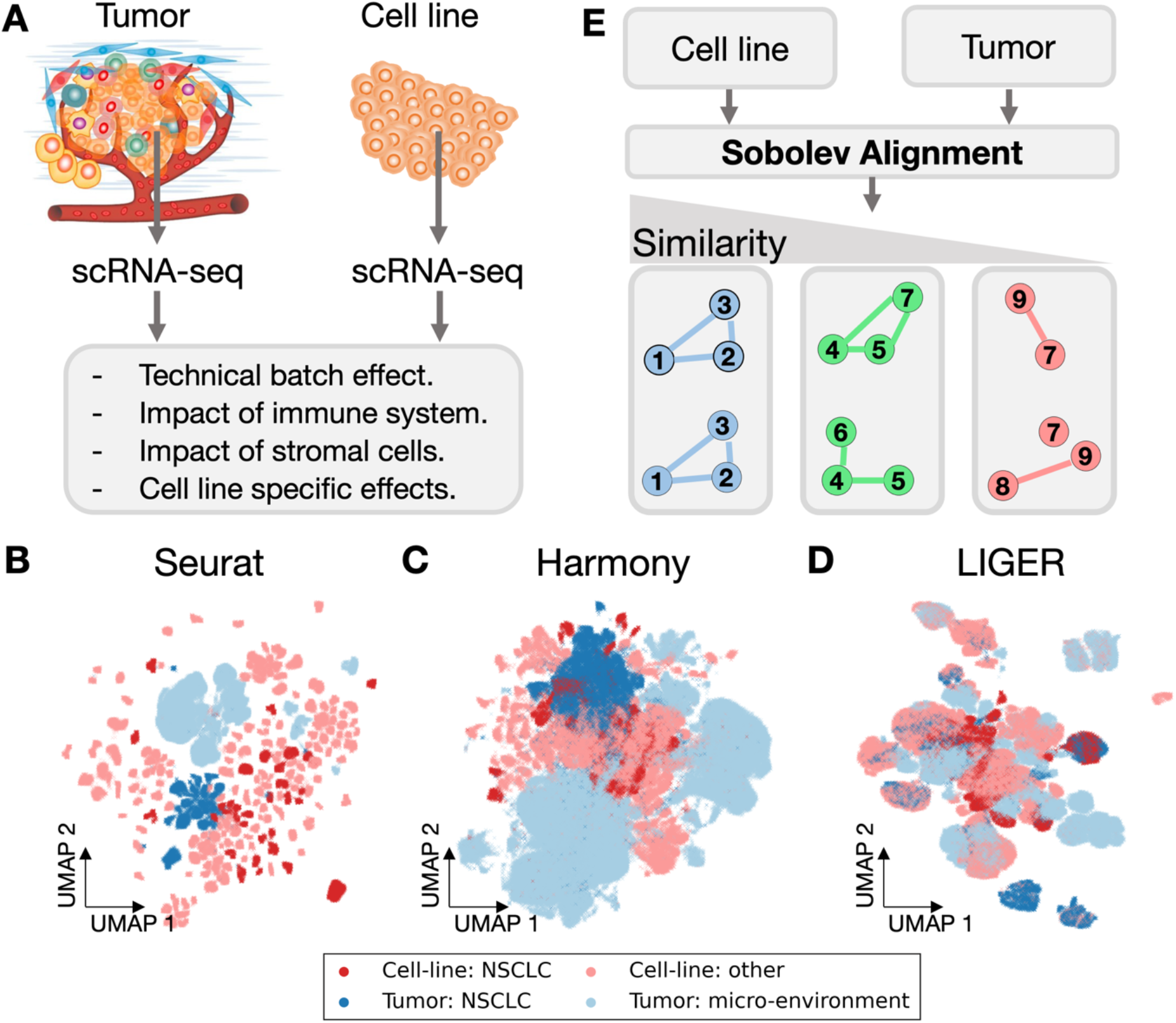
Transcriptional differences between cell lines and tumors obscure the traditional cell type division. (**A**) Schematic model highlighting differences between tumor and cell line gene expression profiles. Although differentiated in the same cell type, cell lines and tumors from the same cancer type are expected to differ in expression. (**B-D**) UMAP projections obtained using three state-of-the-art batch-effect correction approaches: Seurat (**B**) Harmony (**C**) and LIGER (**D**). Cell lines are divided in two cell-type categories: NSCLC (Non-Small Cell Lung Cancer) cell lines in dark red, and other types in light red; patient cells are divided between epithelial cells (coined NSCLC) and other cells (micro-environment) in dark and light blue respectively. (**E**) Diagram of our analysis, which consists in teasing apart biological processes shared between cell lines and tumors from processes specific to either cell lines or tumor.

### Sobolev alignment compares deep probabilistic models by kernel approximation

We previously introduced two computational approaches to compare cell line and tumor gene expression profiles in an unsupervised manner: PRECISE^11^ and TRANSACT^13^, respectively based on PCA and kernel PCA. These two dimensionality reduction methods, however, do not account for the specific properties of scRNA-seq data, such as zero-inflation or over-dispersion, which limit their applicability to single-cell data. To accommodate for these properties, we replaced the (kernel-)PCA dimensionality reduction step with a Variational Auto-Encoder (VAE)^22^ tailored to scRNA-seq data^25,26^. A VAE consists of two neural networks: one network which reduces the dimensionality of gene expression profiles to a much smaller number of *latent factors*, and a second which aims to reconstruct the profiles from the latent factors. Akin to principal components, each latent factor represents a source of variation and captures combinations of correlated non-linear patterns in the studied dataset (**Extended Figure** 2B).

To capture processes present in either cell line or tumor data, we train two independent VAEs (Methods), one on cell lines and one on tumors (**Figure 2**A), instead of a single VAE on the joint cell line and tumor data. This allows each model to capture the variability within either data set separately, while discarding any variability across datasets (**Extended Figure 2**C). Next, based on the similarities between the two resulting sets of latent factors, cell line latent factors are matched with tumor latent factors based on a cosine similarity score (**Figure 2**B, Methods), where a high cosine similarity score indicates that two factors are overlapping and thus share underlying biology. Note that this cosine similarity is determined at the gene-level and takes into account how each gene, as well as each nonlinear combination of genes, influences the latent factor. The cosine similarity matrix between the two sets of latent factors is then used to generate pairs of matching factors (one from the cell lines and one from the tumors) that are ordered by decreasing similarity following the notion of Principal Vectors^37^ (**Figure 2**C, Supp. Methods). Consequently, each resulting *Sobolev Principal Vector* (SPV) corresponds to a pair of processes – one from the cell line and one from the tumor data. By construction, the first SPVs correspond to non-linear combinations of genes similar in cell lines and tumors, whereas the later SPVs relate to cell line or tumor-specific biology. We show that the SPVs can be interpreted by exploiting a closed-form solution for the Taylor expansion of the Gaussian kernel^38^ (**Figure 2**E), which computes the contributions of genes and their interaction terms to each SPV (Methods). The gene-contributions are analyzed using standard PreRanked GSEA^39^, while we derived an extension of GSEA for the interaction terms (Methods). Sobolev Alignment relies on several hyper-parameters, which we selected as indicated in **Extended Figure 2**A.

**Figure 2.**
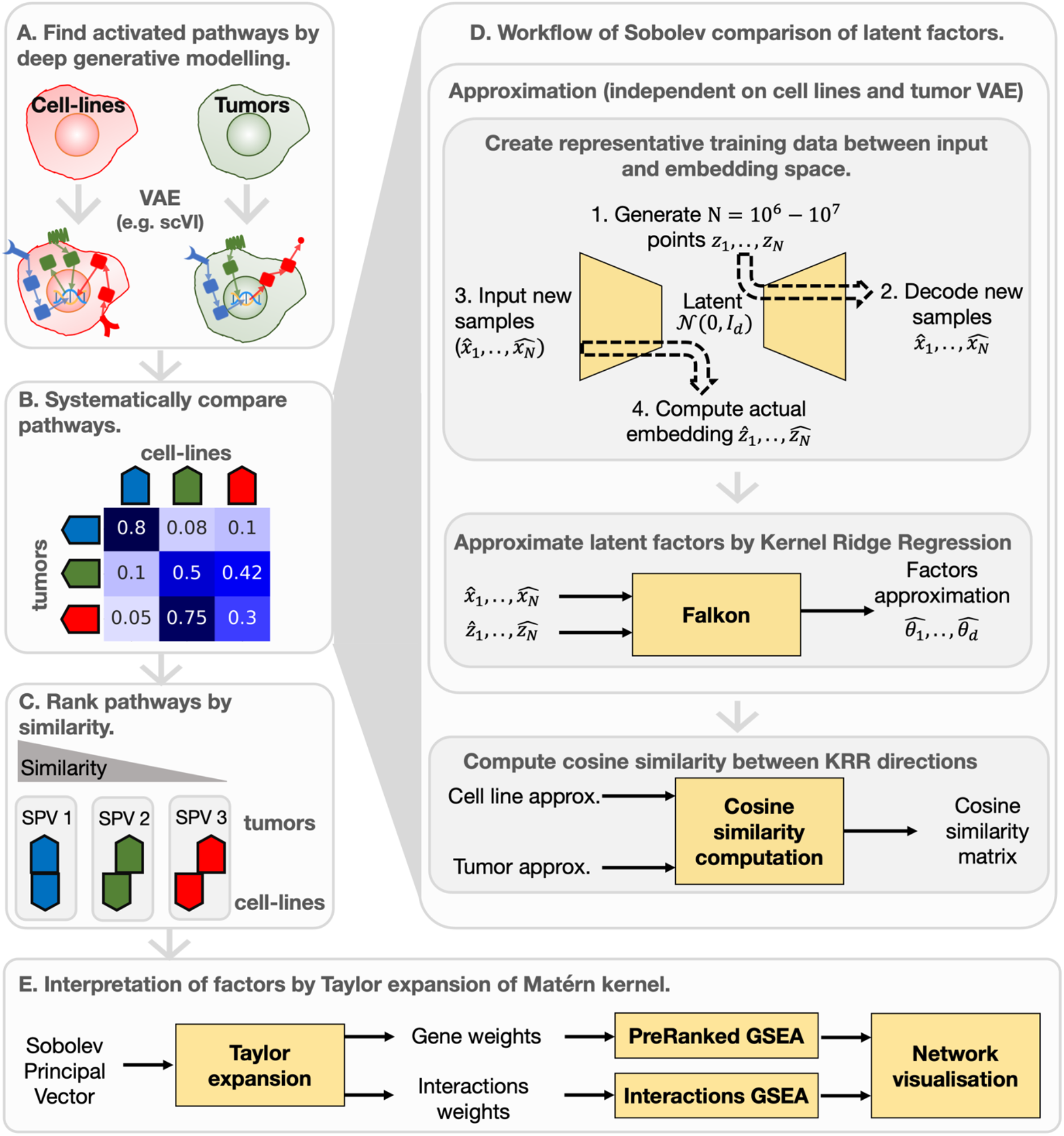
Sobolev Alignment systematically compares molecular processes active in cell lines and tumors. (**A**) In the first step, we employ Variational Auto-Encoders (VAE) to aggregate single cell profiles into a handful of so-called *latent factors*. Each of these latent factors represents a complex non-linear gene combination. Leveraging a model tailored to scRNA-seq data, scVI, this decomposition also accounts for technical issues, such as dropout, dispersion, or library-size. We train two independent VAEs, one for each data stream, with different architectures (Methods, **Supp. Figure** 2A), resulting in two sets of latent factors. (**B**) In order to relate these two sets of factors, Sobolev Alignment yields a cosine similarity matrix with values ranging from 0 to 1: 0 means that the biology supporting the two factors is completely different while 1 means that the genes supporting the latent factors are perfectly similar. (**C**) The cosine similarity matrix is finally decomposed to obtain Sobolev Principal Vectors (SPV) which are pairs of latent factors – one from cell lines (top), one from tumors (bottom) – ranked by gene-level similarity. Here, the first SPV corresponds to two highly similar processes, while subsequent SPV pairs contain less and less similar processes. (**D**) Technically, Sobolev Alignment leverages the generative nature of each VAE to build a large labeled training dataset. This dataset is then employed as labeled training data in a large-scale Kernel Ridge Regression, using Falkon, which approximates the encoder functions. We approximate cell line and tumor VAEs by two different sets of kernel machines, independently from one another but using the same Matérn kernel (Methods). We then compare these two sets of approximations, yielding the cosine similarity matrix (Supp. Methods). (**E**) In order to interpret the SPVs, we derived a scheme which relies on the Taylor expansion of the Gaussian kernel. This yields two sets of contributions: the contribution of genes and of interaction terms — which are the products of two genes. These weights are used in a Gene Set Enrichment Analysis (GSEA) framework to measure the enrichment of certain biological processes.

In the proposed procedure there is one limitation: the latent factors learned by the VAE are difficult to interpret and there is no simple manipulation to obtain the contributions of the genes to the factors. We propose to overcome this by approximating the VAE mapping using a Matérn kernel machine (**Figure 2**D), able to approximate a very wide class of functions, so-called Sobolev spaces, and mathematically easy to handle (Supp. Material). Note that we cannot initially use this kernel machine to find the latent factors as they assume a gaussian noise model which is violated in scRNA-seq data. As the VAE is a generative model, we can train the kernel machine by generating a large data set (~10^6^-10^7^ data points) from the trained scVI models (**Figure 2**D). Each latent factor is then approximated using Falkon^23,24^, a large-scale Kernel Ridge Regression tool based on the Nyström approximation (Methods).

### Synthetic data comparison

In order to demonstrate the utility of SA, we applied it to synthetically constructed single cell datasets. We employed Dyngen^40^ to generate the single cell data using three synthetic models. In each model we simulated processes associated with the source (cell line) and a target (tumor), respectively, and we varied the level of overlap (conservation) between the source and target.

For the first model (*model I*) the source and target consist of a conserved regulatory network (A1-A4), which is complemented with another (independent) network; X1-X4 for the source and Y1-Y4 for the target (**Figure 3**A). Following *Cannoodt et al*, each network is initialized by “external” genes (Ext1-Ext6) (**Figure 3**A). Following Dyngen, source and target datasets furthermore differ by their kinetics parameters and the structure of housekeeping genes and transcription factors, which are used for generating the data, but excluded when applying SA. We ran the complete SA pipeline (**Extended Figure 2**A) and obtained 8 SPVs for source and 10 for target. To inspect the SPVs, we first look at their linear portion (**Figure 3**B), which explains 50% of the SPVs (**Extended Figure 3**B-C). The three top source and target SPVs are exclusively made of signal in the conserved network. The subsequent two SPVs are made of a mix of conserved and source/target-specific signal. Finally, the last three SPVs are made solely of source/target-specific genes. Next, we look at the interaction genes (**Figure 3**C), which explain 30% of the SPVs (**Extended Figure 3**B-C) and observe a similar pattern as for the linear part. The first 3 SPVs are driven by shared genes (“pure-shared”), while the last 3 SPVs are driven by source/target-specific (“pure-specific”) and hybrid interactions, i.e., involving one shared and one source/target-specific gene.

**Figure 3.**
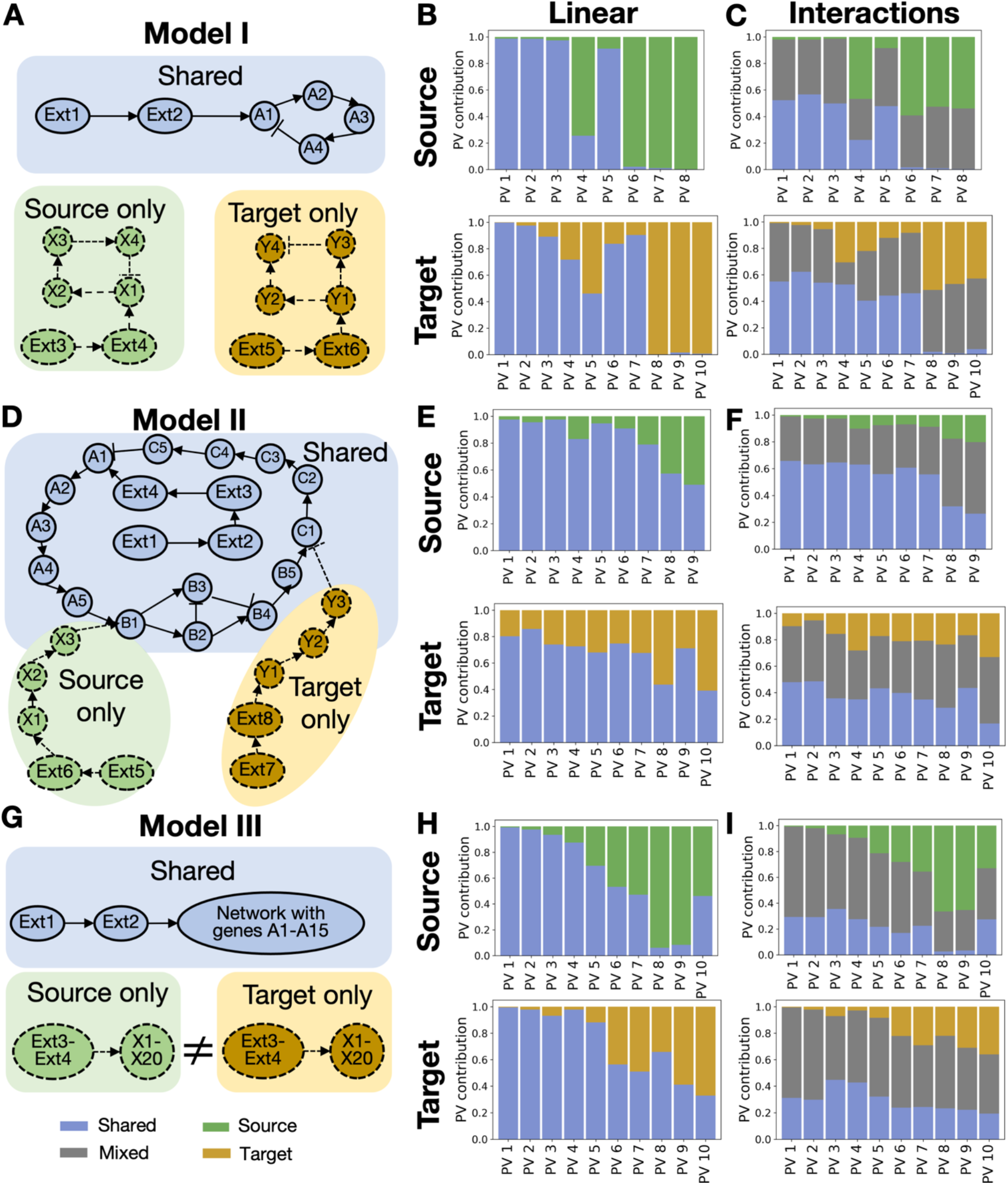
Sobolev Alignment faithfully discriminates shared from specific biological processes on three synthetic datasets. We generated three models of increasing complexity to assess the capacity of Sobolev Alignment in discriminating common from data-specific signals. (**A**) Gene regulatory network of *model I*. A blue box indicates a gene shared between source and target, green a gene specific to source, and orange to target. (**B**) Proportion of gene weights for source (top) and target (bottom) PVs. (**C**) Proportion of interaction weights for source (top) and target (bottom) PVs. (**D-F**) Results similarly displayed for *model II*. (**G-I**) Results similarly displayed for *model III*. (blue: common; grey: interactions with one common and one specific gene; green: source-specific; orange: target-specific)

Next, we constructed a more challenging dataset with a conserved negative feedback-loops (like, for example, mitotic pathways) and source/target-specific external control. Following an example proposed by *Cannoodt et al*, we designed a conserved cycling backbone pathway (**Figure 3**D). Taking inspiration from the MAPK pathway, we added a linear pathway branching from an external gene (Ext-5) to a cell-cycle gene (B1) for the source dataset. Taking inspiration from DNA damage repair pathways, we added a linear pathway inhibiting the end of the cycle (C1) for the target dataset (**Figure 3**D). Analysis of the linear part (**Figure 3**E) and the non-linear part of the SPVs (**Figure 3**F) showed a negative trend of shared linear and interaction terms. Shared linear terms account for 98% of source SPV 1 and 80% of target SPV 1, while these proportions decrease to 50% and 40%, respectively, for SPV 9. A similar trend is observed for the interaction terms. Note that none of the SPVs are solely determined by conserved signal or source/target-specific signal (as was the case for *model I*). This might be because, for *model II*, the individual genes have a direct influence on the conserved network. As a result, a significant amount of covariance is to be expected between the shared and source/target-specific genes. A factor model such as scVI would then also be unable to disentangle the different origins and create factors which mix both individual and common genes (**Extended Figure 4**). The SPVs resulting from SA do, however, capture a ranking of components according to the involvement of the conserved part between the source and target datasets.

Finally, models I and II harbor individual genes that are specific to either source or target, resulting in genes clearly over-expressed in their specific model. To further challenge our Sobolev Alignment, we created a model (*model III*, **Figure 3**G) that has a conserved feedback-loop similar as in *model II* (genes A1-A15), and a set of conserved genes X1-X30 but whose wiring is different between the source and target. When applying SA, we observed that the linear part of the first three SPVs are solely made of shared genes (**Figure 3**H) and that the non-linear part of these SPVs is dominated by either pure-shared or hybrid interactions (**Figure 3**I). Also here, the last three SPVs have a limited contribution of shared genes or interaction terms consisting of conserved genes. The results from these three models demonstrates that Sobolev Alignment is able to capture the shared information between two datasets in the top ranked SPV components.

### Sobolev Alignment effectively integrates cell lines and tumors

We applied our Sobolev Alignment to compare treatment naïve NSCLC cell lines (*Kinker* dataset) and epithelial tumor cells from NSCLC patients (*Kim* dataset) (Methods). Following the hyper-parameter selection pipeline (**Extended Figure** 2), we selected two different architectures for the scVI models (**Supp. Table** 1) and approximated the embedding of the VAE latent factors with the Falkon kernel machine using a Laplacian kernel with *σ* = 15 (Methods). As our alignment relies on how well each scVI latent factor is approximated, we first confirmed the goodness of fit of our approximation. For each factor, we calculated the Spearman correlation between our approximation and the scVI-computed values (**Extended Figure 7**A). We observed that the cell line factors are approximated with a Spearman correlation between 0.97 and 0.99, while the tumor factors are approximated with a Spearman correlation ranging from 0.95 to 0.99, both indicating that the kernel machine faithfully approximates the VAE embeddings.

The similarities between the SPVs of the cell lines and the tumor data ranged from 0.51 to 0.01 (**Extended Figure 7**B). To only retain significantly similar pairs of SPVs, we compared the obtained similarities against a null model (**Extended Figure 7**B). This resulted in 12 SPVs that we employed for further analyses. From the 12 cell line-tumor SPV pairs, we subsequently constructed a 12-dimensional consensus space (Sobolov Consensus Space, SCS). Each dimension of the consensus space consists of a consensus vector obtained by interpolating between the matched cell line and tumor SPV pair (Methods). These consensus vectors represent the best balance between the effects of the cell lines and the tumors. We then projected the cell line and tumor data on the resulting 12 consensus vectors and performed a UMAP projection and observe a continuity between cell line and tumor clusters but no co-clustering (**Figure 4**A). Inspired by Seurat’s and LIGER’s workflows, we performed a Mutual Nearest Neighbor (MNN) correction to the cell-line and tumor datasets on Sobolev Consensus Space. Although MNN on the gene expression profiles performs poorly (**Figure 4**B), its combination with SA resulted in a clear improved co-clustering of cell lines and tumors (**Figure 4**C). Such co-clustering is not achieved by either Seurat v3 (**Extended Figure 8**A), Harmony (**Extended Figure 8**B), or LIGER (**Extended Figure 8**C). In subsequent analyses, we will employ the datasets resulting from the projection on the Sobolev Consensus Space with MNN correction, denoted *SA+MNN* space.

**Figure 4.**
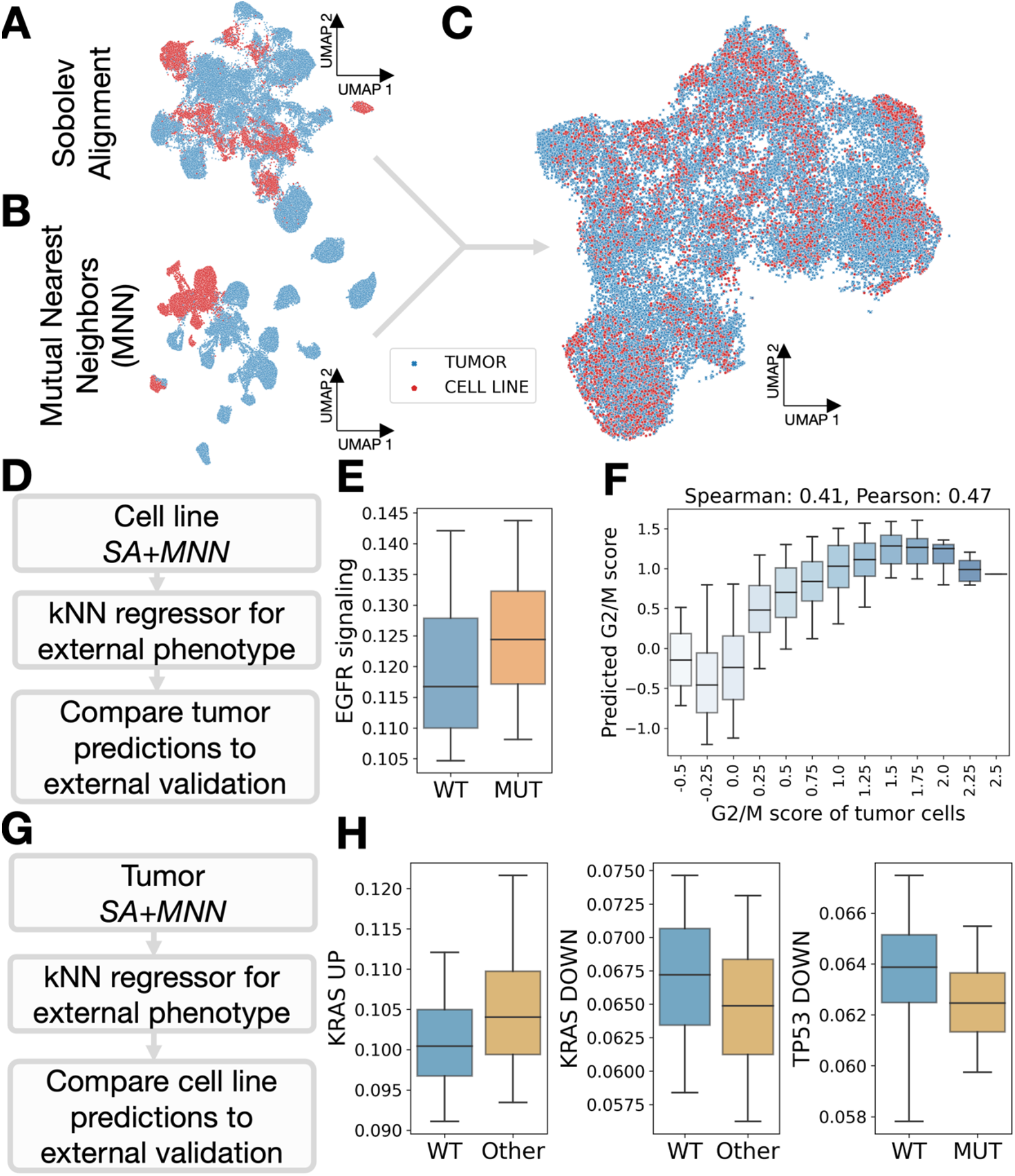
Sobolev Alignment effectively integrates cell lines and epithelial tumor cells. We employed Sobolev Alignment to compare untreated epithelial cells from NSCLC cancer patients with untreated NSCLC cell lines. (**A**) UMAP of cell lines and tumors after projection on SPVs and interpolation. (**B**) UMAP of cell lines and tumors after correction with Mutual Nearest Neighbors (MNN). (**C**) UMAP of cell lines and tumors after projection on SPVs, interpolation, and subsequent correction by MNN. (**D**) Workflow of our cell line neighborhood validation, which computes, for each tumor cell, the value of a certain phenotype in neighboring cell line cells. (**E**) Relationship between imputed EGFR signaling levels and EGFR status in tumor cells. (**F**) Relationship in tumor cells between G2/M score imputed from cell lines (y-axis) and G2/M score computed using Seurat v3 (x-axis). (**G**) Workflow of our tumor neighborhood validation, reversing the order from cell line neighborhood validation. (**H**) Relationship between pathway levels and oncogenic form in cell line; left to right: genes up-regulated in KRAS mutated cells (“KRAS UP”), genes down-regulated in KRAS mutated cells (“KRAS DOWN”) and genes down-regulated in TP53 mutated cells (“TP53 DOWN”) (**Extended Figure 9**H-M).

To validate the quality of our alignment, we compared the EGFR mutation status of patients from the *Kim* (tumor) dataset with the EGFR signaling activity in the cell lines; cell lines in the *Kinker* dataset do not harbor any activating EGFR mutation, rendering impossible the direct transfer of mutation status. The cell line EGFR signaling is computed^41^, for each cell, using UCell^42^ (Methods). As we expect cell line cells with high EGFR signaling activity to be in close proximity to EGFR mutant tumor cells, we predict, for each tumor cell, the EGFR signaling level from the cell line using a k-Nearest Neighbor (kNN) regression in the SA+MNN space (**Figure 4**D, **Extended Figure 9**A). From the cell-line predictions, we observed that wild-type EGFR tumors have the lowest EGFR signaling level (**Figure 4**E, **Extended Figure 9**B), coherent with a constitutive activation of EGFR signaling pathway. In other words, EGFR-mutated tumor cells indeed preferentially cluster with cell lines harboring a high level of EGFR signaling after aligning the datasets to the *SA+MN* consensus space. Next, we evaluated whether the cell cycle states of neighboring cell line and tumor cells in the SA+MNN space are comparable. “G2/M” and “G1/S” scores were provided for the *Kinker* cell line dataset. We computed equivalent quantities on the *Kim* tumor dataset using the Seurat v3 cell-cycle regression tool. Comparing the tumor cells’ G2/M scores predicted with a kNN regression model from the cell line data (**Extended Figure 9**C) with the cell-cycle scores, we observe a spearman correlation of 0.41 (**Figure 4**F), indicating that our co-clustering captures mitotic entrance. When performing the same experiment with the S scores (**Extended Figure 9**D-E), we however only observe a modest spearman correlation of 0.11. Following a similar type of analysis (**Figure 4**G), we find that tumor cells in the neighborhood of KRAS-mutated cell lines have a higher level of genes over-expressed in KRAS-mutant lung cancer cells (“KRAS UP”) and a lower level of genes of genes down-regulated in KRAS-mutant lung cancer cells (“KRAS DOWN”) (**Figure 4**H, **Extended Figure 9**F-I). Also, tumor cells in the neighborhood of TP53 mutated cell lines have lower levels of genes down-regulated by TP53 mutations (**Figure 4**H, **Extended Figure 9**J); however, we could not find an association for genes up-regulated by TP53 mutation (**Extended Figure 9**K-L). Taken together, alignment of cells from cell lines and tumors into the *SA+MNN* space conserves important biomarkers, indicating the biological relevance of our mathematical model.

### Sobolev Alignment highlights the conservation of important intrinsic immune-related pathways in cell lines

To assess which tumor-related biological processes align well to cell line biology, we analyzed each of the tumor SPVs (**Figure 4**). First, by construction, cell line and tumor cells in the same neighborhood show similar values for all SPVs, indicating that SPVs can be understood as biomarkers, which can help relate clinical samples to cell line models. To interpret the tumor SPVs, we computed the linear (Methods, **Supp. Table** 2) and interaction loadings (Methods, **Supp. Table 3**), and subsequently performed a gene set enrichment analysis (Methods, **Supp. Table 4**). Using GSEA on the linear part (**Figure 2**E), we report, for each SPV, the top 10 most enriched gene sets in the linear portion (FDR < 0.05). To study interaction terms, we created interaction-gene-sets that correspond to a pair of gene sets (Methods). For instance, the interaction-gene-set “M Phase x Keratinization” contains all interaction pairs with one gene from the “M Phase” gene set, and one gene from the “Keratinization” gene set. For each SPV, we computed the Normalized Enrichment Scores (NES) of all interaction gene sets using a modified procedure akin to PreRanked GSEA (Methods). The remaining analyses are based on the top 5% interaction terms with largest NESs in absolute value.

We observed that negative coefficients of SPV 1 are enriched for gene sets linked to G2/M mitotic gene sets (**Figure 5**A), indicating a conservation of mitosis-related biological processes. Keratin genes are also driving the enrichment of several gene sets in the negative coefficients (e.g., keratinization). Four known lung adenocarcinoma markers^43^, KRT7-8–18-19, are amongst the 10 most important linear contributors, indicating a conservation of the keratin markers between cell lines and tumors. The two subsequent SPVs (SPV 2 and SPV 3) are enriched for PI3K-AKT and the MET pathways (**Figure 5**B-C), two important cancer-related pathways expected to be shared between cell lines and tumors. SPV 4 is not associated with any enrichment. Positive coefficients of SPV 5 (**Figure 5**D) are significantly enriched for immune-related gene sets (interferon signaling, cytokine signaling, interferon gamma, adaptive immune system) and their interactions. The negative coefficients of SPV 5 are enriched for keratin-related gene sets. Analysis with the oncogenic signatures from MSigDB (**Supp Table 4**) indicates an enrichment in the positive portion for genes up regulated in KRAS mutant cells (FDR < 0.035). SPV 5 values gradually change across the UMAP, suggesting a higher-level structure shared between cell lines and tumors. More specifically, cells in the lower-left of the UMAP exhibit a high keratinization level, while cells in the upper-right present either KRAS activation, or a high level of activity for immune-related pathways. Analysis of SPV 6 (**Figure 5**E) shows a similar pattern as SPV 5 with keratinization and immune activity enriched in the positive and negative coefficients, respectively. SPV 6 also shows a gradient between high-keratinization and high immune-response, focused here on antigen presentation (MHC class II, microbial peptides, antigen presentation). Altogether, these results highlight the conservation of immune-related pathways in lung cancer cell line models alongside their interplay with keratin-levels and KRAS aberrations.

**Figure 5.**
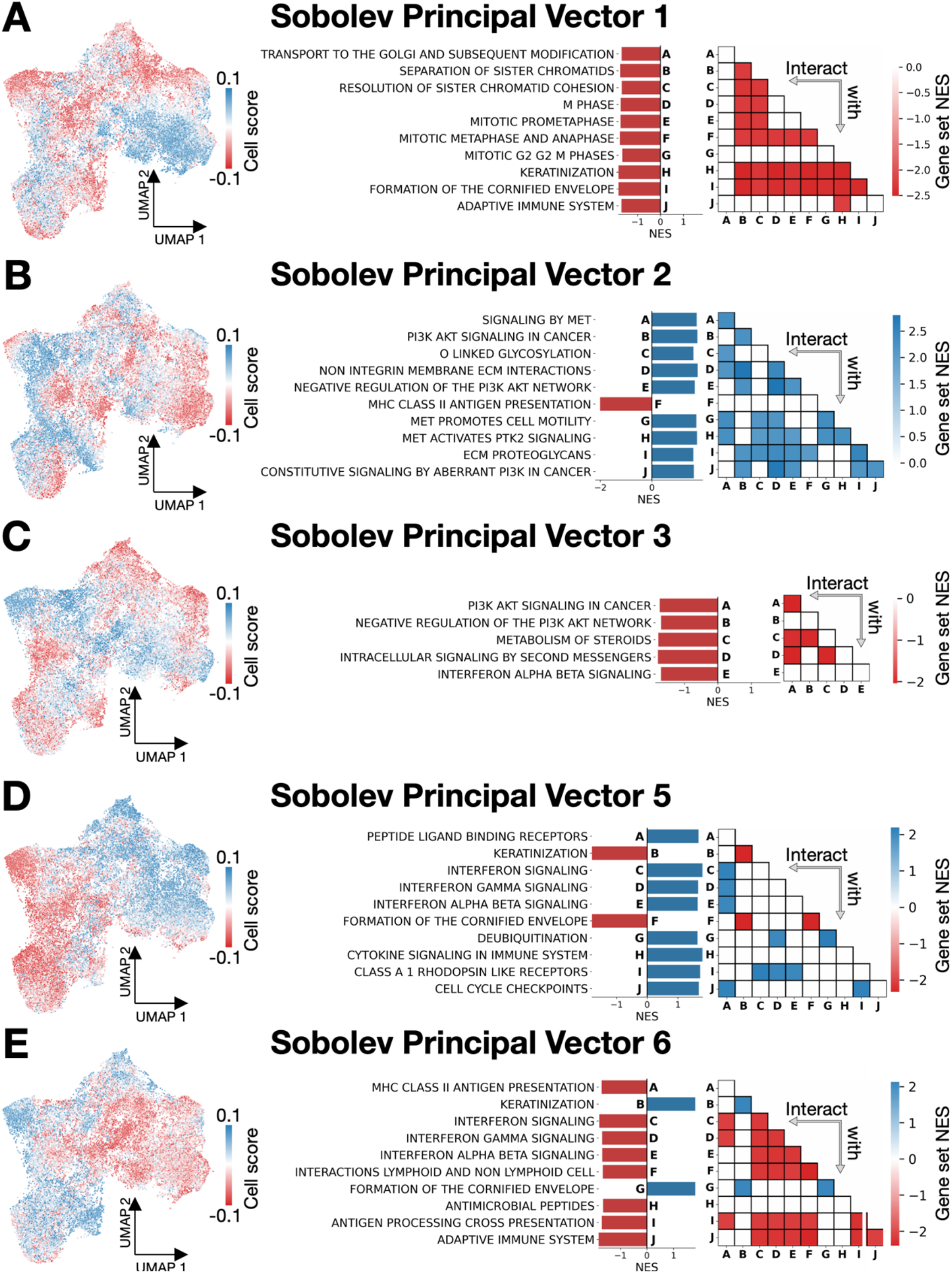
Analysis of Sobolev Alignment directions show the conservation in cell lines of important mitotic pathways alongside innate immune processes. For each SPV computed between the Kinker and the Kim datasets (**Figure 4**), we study the tumor processes recapitulated in cell lines by performing a gene set enrichment analysis on the tumor SPVs. (**A**) Results for SPV 1. The left UMAP of cell lines and tumors is colored by the cell scores of SPV 1. Turning to gene loadings, the subsequent bar plot summarizes the top 10 most enriched gene sets for the Reactome MSigDB collection, represented by their Normalized Enrichment Scores (NES). Finally, interaction terms between these linearly enriched gene sets are reported in a heatmap. A white value indicate that the interaction term is not in the 5% most enriched interaction terms. A negative value (red) indicates that the product of the two pathways drives down the value of the SPV; a positive (blue) value indicates the reverse. The cell scores (left) and gene-loadings (right) are connected: blue cells in the UMAP harbor a high expression of blue linear and interaction gene sets, and reciprocally. We repeated the experiment for SPV 2 (**B**), SPV 3 (**C**), SPV 5 (**D**) and SPV 6 (**E**) with a similar display.

### Sobolev Alignment allows to study the mode of action for certain drugs

Finally, we employed our methodology to study the different modes of action to anti-cancer drugs. We first computed the SPVs between the *McFarland*^19^ cell line and the *Kim* tumor datasets. The *McFarland* dataset consists of a multiplexed perturbation screen on 33 NSCLC cell lines where each cell line was exposed to 19 anti-cancer drugs. Transcriptomic read-outs were measured 6 or 24 hours after drug induction (**Extended Figure** 11). To study this set of perturbations (**Figure 6**A), we first mapped the cell line data to the top SPVs (**Extended Figure 12**A-C). Because tumor cells are now matched with cell line data, drug perturbations on the top SPVs correspond to biological processes corrected for cell-line specificities.

**Figure 6.**
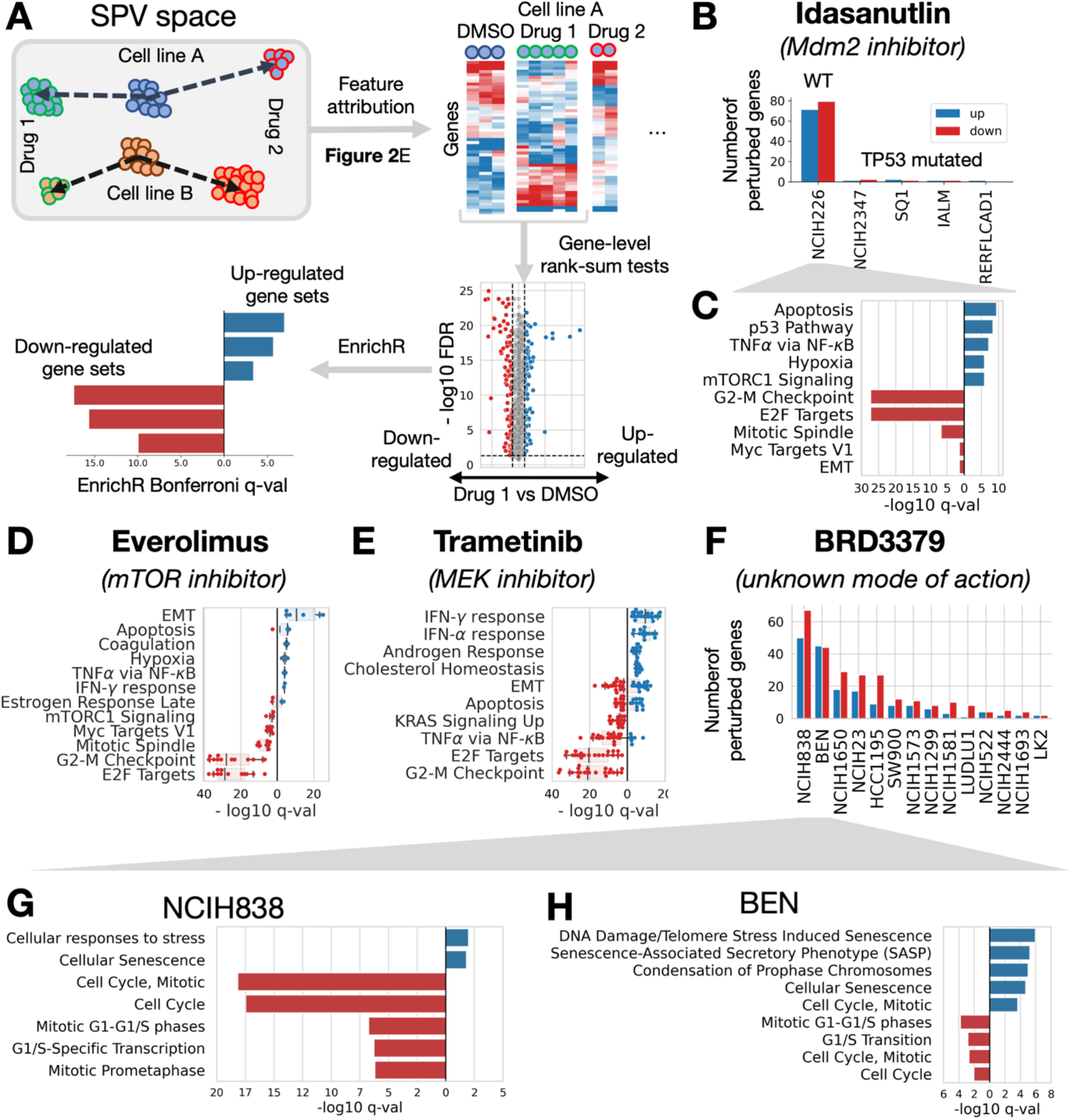
Sobolev Alignment identifies modes of action from drug perturbations. Exploiting a large drug-perturbation screen (McFarland dataset), we studied the modes of action of several drugs recapitulated by the SPV common with tumors (Kim dataset). (**A**) After first embedding all cells to the common SPV space, we computed the gene-weights corresponding to each single cell embedding. Using a rank-sum test (Mann-Whitney, Benjamini-Hochberg FDR multiple-testing correction), we assessed, for each drug, the difference in gene weights between DMSO-treated cells and drug-treated cells; the threshold in effect size was set as the 95% percentile of gene-weights differences observed within the DMSO-treated cells of the studied cell line. Up- and down-regulated genes were then analyzed using EnrichR. (**B**) Number of up- and down-regulated genes for 5 cell lines subjected to Mdm2-inhibitor Idasanutlin alongside their TP53 mutation status. (**C**) Enrichment observed using the MSigDB Hallmarks collection for NCIH226 (blue: enrichment in genes up-regulated by Idasanutlin, red: enrichment in down-regulated genes). (**D**) Boxplot of q-values obtained when analyzing Everolimus (mTOR inhibitor). (**E**) Boxplots of q-values obtained when analyzing Trametinib (MEK-inhibitor). (**F**) Number of up- and down-regulated genes for each cell line perturbed with BRD3379, a drug with an unknown mode of action. Zooming on the two most perturbed cell lines, we performed the EnrichR analysis using the Reactome MSigDB collection for NCIH838 (**G**) and Ben (**H**).

For each embedded cell, we computed the gene-loadings corresponding to the combination of projected SPV scores (Methods). For each cell line and exposed drug, we employed the Mann-Whitney test to assess, per gene, whether the gene weights from cells retrieved after drug induction are significantly different from gene weights of cells obtained after vehicle-treatment (DMSO). To include a measure of effect size, we filtered genes based on the mean differences of associated gene-weights between the two groups (drug induction and DMSO). The threshold is set to the 95% percentile of this effect size computed by comparing random DMSO-perturbed cells (95% percentile) (**Extended Figure 13**A). Genes which are significant (FDR < 0.05) and pass the effect size filter are considered to be perturbed by the drug (for the cell line of consideration). The set of genes is then further annotated using EnrichR^44^ to associate them with biological processes (only gene sets with at least 5 perturbed genes are considered).

We first analyzed Idasanutlin (a.k.a. Nutlin-3), an Mdm2 inhibitor which triggers apoptosis selectively in TP53 wild-type cells^45^. When examining the number of genes perturbed after 24 hours for each of the 5 cell lines (**Figure 6**B), we observe a clear difference between the sole TP53 wild-type cell-line (NCIH226) and the four mutated cell lines. While NCIH226 harbors 70 up-regulated genes and 78 down-regulated genes, none of the mutated cell lines show more than 5 perturbed genes. For NCIH226, we observe a clear enrichment for apoptosis and p53 pathway in the up-regulated genes, while G2M checkpoints and E2F targets are down-regulated, coherent with the known mode of action of Idasanutlin (**Figure 6**C). We then turned to Everolimus, an mTOR inhibitor^46^, and performed the analysis presented in **Figure 6**A for each of the 18 perturbed cell lines (**Extended Figure 13**B, Supp. Table). The enrichment scores are summarized in a boxplot (**Figure 6**D), each point corresponding to a significantly enriched cell line. We observe that for 12 cell lines, G2M checkpoints and E2F targets are significantly downregulated; the 6 other cell lines correspond to the cell lines with the least number of perturbed genes (**Extended Figure 13**B). We also observe a down-regulation of mTORC1 signaling for four cell lines (A549, IALM, LU99, NCIH1355), coherent with the known mode of action. We performed a similar experiment for Trametinib, a MEK-inhibitor (**Figure 6**E, **Extended Figure 13**C). Upregulation of IFN*γ* and IFN*α*, downregulation of genes over-expressed in KRAS mutant cells, and downregulation of cell cycle pathways are observed, consistent with previous reports^19,47^. Androgen response and cholesterol homeostasis pathways are enriched, suggesting combination therapies with anti-androgen or cholesterol lowering drugs may yield synergistic effects or reduce resistance to MEK inhibitors^48–50^. Several pathways, such as Apoptosis and EMT, display a duality where up-regulation is observed for some cell lines while others show depletion. While this may represent normal variations in drug response, these results are consistent with the notion that resistance to MEK inhibitors can be achieved through various feedback mechanisms, including EMT, and highlights the need for combination therapies in NSCLC^51,52^. Finally, we turned to a drug with an unknown mode of action, BRD3379. When examining the number of perturbed genes per cell line, we observe that two cell lines stand out: NCIH838 and BEN (**Figure 6**F, **Extended Figure 13**D). Performing pathway enrichment analysis revealed that both NCIH838 (**Figure 6**G) and BEN (**Figure 6**H) are characterized by cell cycle arrest (down-regulation of cell cycle gene sets). Both cell lines show enrichment for genes involved in senescence, reminiscent of the effects of BRD4 inhibition^53,54^. Indeed, genes shown to be related to senescence induction by BRD4 inhibition, such as Aurora kinases A/B^55^ and CDKN1A (p21)^54^, show similar patterns of expression upon BRD3379 treatment (**Supp. Table** 5). Furthermore, JQ1 inhibition similarly shows induction of senescence and cell cycle arrest (**Extended Figure 13**E-F). Therefore, we posit that BRD3379 mode of action is likely to be similar to that of BRD4 inhibitors. Taken together, these results demonstrate the ability of our approach to recapitulate clinically validated drug response mechanisms and exemplify the potential of Sobolev Alignment to decipher complex modes of action.

## Discussion

We showed that single cell profiles measured from pre-clinical models and human tumors cannot be aligned using current batch-effect correction tools like Seurat, LIGER and Harmony. To help researchers study the translational potential of pre-clinical models, we derived a novel framework that exploits the power of unsupervised deep generative models (VAE), known to their ability to embed scRNA-seq data, with the benefits of kernel machines, known to be interpretable. In the first step, a VAE is trained to embed both input datasets independently, capturing the different sources of variations into a set of latent factors. In the second step, we approximate the mapping towards the latent factors using Falkon-trained kernel machines, which allows us to calculate the contribution of each gene to each of the latent factors. We then match the latent factors of the two domains by calculating their Sobolev Principal Vectors (SPVs). Finally, we construct a consensus space by interpolation between matched SPVs onto which all data can be projected.

We applied our approach to a set of synthetic examples and showed that a conserved network of genes contributes to the most important SPVs. We then applied our alignment to NSCLC cell lines and epithelial cancer cells and showed enrichment of known oncogenes in the top SPVs. Further analysis of the top SPVs pointed towards the conservation of mitotic and immune-related pathways in cultured cells. In a last experiment, we aligned a multiplexed drug-perturbation screen with NSCLC cell lines and showed that the common SPVs recapitulate modes of action for several drugs. Although we restricted our analysis to single cell gene expression data, our computational approach is versatile, and can easily be adapted for other molecular features, such as chromatin accessibility^56^, protein levels^57^ or ribosomal profiling^58^. Recently developed deep probabilistic models tailored for such data^59,60^ could be for instance employed to adapt our framework. We also solely compare cell line models to human tumors, but more complex models could be studied equally well, such as organoids or patient derived xenografts.

On a more technical note, our approach develops and exploits connections between deep-learning-based algorithms and kernel methods^61^. Recent works have shown theoretical connections^62^, demonstrating, for instance, the equivalence between the Laplacian kernel and the so-called Neural Tangent Kernel^63^. We envision that future theoretical works could help improve our approach, and we believe that a promising avenue for improvement lies in replacing the kernel ridge regression step. Indeed, we observe that kernel ridge regression is locally adaptive and is not ideal for approximating functions that exhibit large variations in localized areas, as is often the case with neural networks. Furthermore, our interpretation scheme relies on the decomposition of the Gaussian kernel, which we extended to the Laplacian kernel by exploiting connections between the feature spaces of Gaussian and Laplacian kernels. Derivation of a closed-form solution for the Laplacian kernel RKHS basis could help improve our analysis. Finally, our approach could be used in other areas, for instance to define a similarity measure between neural networks, as spearheaded by recent works^64–66^.

## Methods

### Data download and processing

Cell line and tumor data were downloaded following the protocol indicated in the supported publications^18,20^, i.e., using the Broad Institute Single Cell Portal for cell lines, and GEO repository GSE131907 for tumors. Both datasets were processed using Scanpy^67^ (Supp Material).

### Batch effect correction

The Seurat R implementation (version 4.0.1) was used and corrected for batch effect using Reciprocal PCA. We used Liger R implementation (version 0.5.0.9). Finally, we use Harmony Python implementation (version 0.0.5). For all methods, we used default parameters.

### Model selection for scVI

The first step of Sobolev Alignment consists in training two Variational Auto-Encoders (VAEs): one using the source (cell lines) and one using the target (tumor) dataset. For training these two VAEs, we turned to an established implementation, called scVI^26,68^. As any neural-network-based approach, scVI requires a lot of hyper-parameters to be tuned, e.g., number of hidden layers, number of latent factors, number of neurons per layers, dropout rate, weight decay, likelihood function, early stopping or number of iterations. In order to select a combination of hyper-parameters that best reconstructs the data, we turned to Bayesian Optimization^69^ and selected independently for each dataset the best combination of hyper-parameters based on the reconstruction error computed on an held-out test set (Supp. Table 1). The two resulting optimal models were then trained using the complete source and target data, respectively. Furthermore, VAEs are frequently suffering from posterior collapse^70,71^, i.e., the presence of at least one latent factor with zero variance, which causes matrices 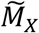 and 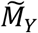 to be singular and renders our alignment strategy unstable (Eq. (5)). To avoid it, we devised a rejection scheme: after complete training of the model, we computed the variance of each latent vector and restarted the training, should one latent vector happen to be collapsed. If the training fails five times in a row, one latent factor is removed.

Naming *d_X_* and *d_Y_* the number of latent factors for source and target, respectively, this training step generates to set of encoders: 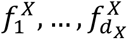 for source and 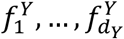 for the target. Each of these functions maps a single cell expression profile to a latent factor score.

### Approximation of latent factors embedding function by Kernel Ridge Regression

In a second step, we approximate each embedding function (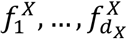 and 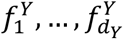) using Kernel Ridge Regression (KRR), with a view to exploit interesting mathematical properties of kernel methods (Supp. Sect. 2). To generate enough training data, we exploit the generative nature of the two scVI models. We provide here the procedure for the source model, which can be readily applied to the target model as well (Supp. Sect. 4).

First, we randomly sample *N* random noise vectors 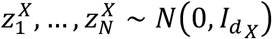, with *I_d_X__* the identity matrix of size *d_X_*. Using the source model decoder, we collect *N* artificial gene expression profiles 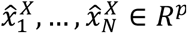. As VAEs are not bijective, the original vectors 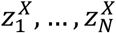 do not perfectly equate the encoder outputs for 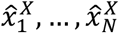. To limit the noise in our model, we input the samples 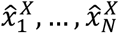 in the encoder functions, resulting in new artificial vectors 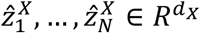; each 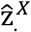 corresponds here to the mean of the embedding parametrization of a sample. We subsequently approximate the embedding functions by training, on the artificial data 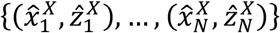, *d_X_* independent KRR models. To scale to the large amount of data we sampled, we turned to Falkon^24^, a stochastic approximation of KRR which relies on the Nyström approximation^72^. Critically, Falkon relies on the choice of a kernel function, which in our case, must provide a universal approximation. We turned to the Laplacian, Matérn and Gaussian kernels, which are defined as follows:

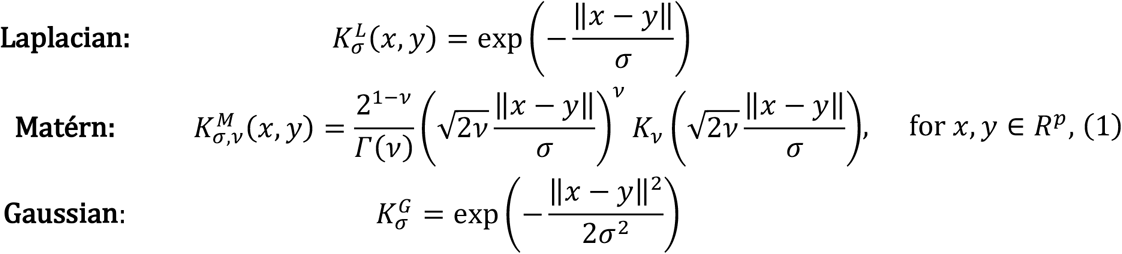

with *σ*, *v* > 0, *Γ* the gamma function and *K_v_* the modified Bessel function of the second kind of order *v*. The parameter *σ* represents the width of the kernel (also referred to as length-scale), while the parameter *v* represents the smoothness parameter (Supp. Sect. 2.2 and 2.3). Let us denote by *K* the kernel selected (Eq. (1)). Falkon selects *M* anchor points (*M* ≪ *N*) among the training data, denoted 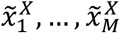 and approximates the embedding functions 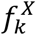 by computing the matrix *α^X^* ∈ *R^d_X_×M^* such that:

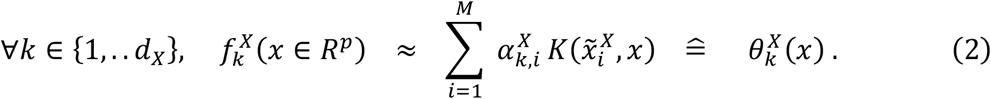

Eq. (2) corresponds to the regular basis-expansion form resulting from the Representer Theorem, with the *M* anchor points instead of the *N* training samples. It is however important to note that the coefficients *α^X^* are optimized using the *N* – *M* non-anchor points, and therefore exploits the whole dataset.

### Comparison of latent factors by Sobolev alignment

We have approximated the source encoder functions 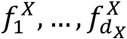 by the kernel machines 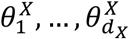, and the target encoder functions 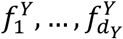 by the kernel machines 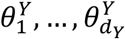. These two approximations are then used to compute the cosine similarity matrix *M* (Supp. Def. 4.10). We here present the computational approach to compute *M*, and refer the reader to Supp. Sect. 4 for the precise mathematical definition and derivation.

Let us define the three following kernel matrices:

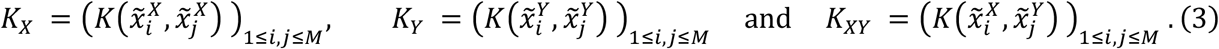

These three matrices correspond to similarity (or kernel) values between the different anchor points computed by Falkon (Eq. (2)). We then define the three following matrices, referred to as un-normalized cosine similarity matrices (Supp. Prop. 4.9.):

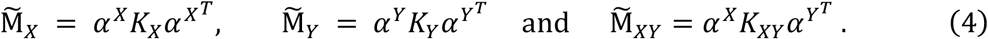

The cosine similarity is then computed as follows (Supp. Def. 4.10.):

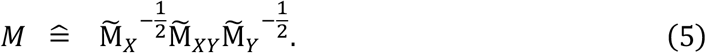

### Computation of principal vectors and principal angles

We now have two sets of vectors which approximate the two VAEs: 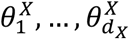 for source and 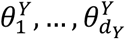 for target. To compare them, we use the notion of Sobolev Principal Vectors (SPVs) which correspond to pairs of vectors (one from source, one from target) ranked by decreasing similarity (Supp. Def. 5.1.). To compute them, we need to decompose the cosine similarity matrix (Eq. (5)) by SVD:

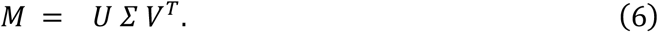

The diagonal matrix *Σ* contains the similarity values between the principal vectors (Supp. Cor. 5.2.1.), and can be written as:

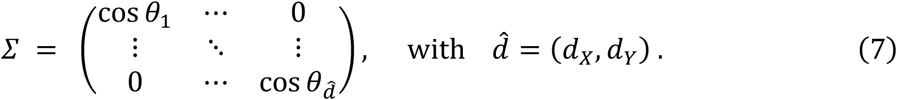

We also define the two matrices 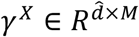 and 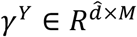 as

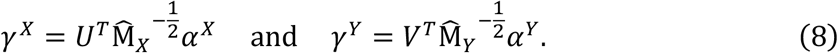

The principal vector pairs 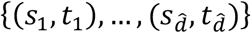 are finally computed as (Supp. Theorem 5.2.):

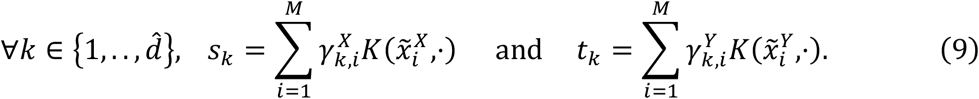

### Interpretability of principal vectors by kernel Taylor expansion

We here present the interpretability scheme which relies on the explicit characterization of the Gaussian kernel RKHS^39^ (Supp. Sect. 6). We define the following *univariate basis function* 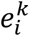, for *i* ∈ {1,…, *p*} and *k* ∈ *N*, as:

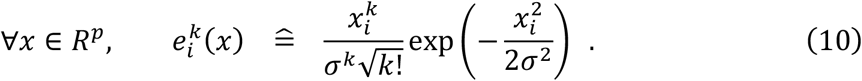

An orthonormal basis of the Gaussian RKHS can then be defined by combining these univariate basis functions with an outer product (Supp. Prop. 6.3), yielding *Gaussian basis functions*, defined, for *K* = [*k*_1_,…, *k_p_*] ∈ *N^p^*, as:

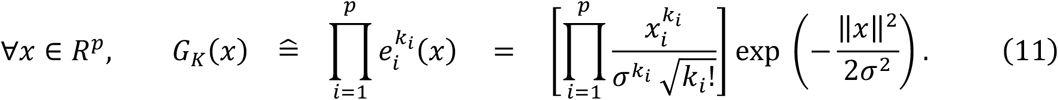

Since these Gaussian basis functions form an orthonormal basis, we exploit them to parametrize the latent factors and the SPVs (Supp. Subsect. 6.1). Based on Eq. (11), we define the following offset matrices as the exponential term, i.e.,

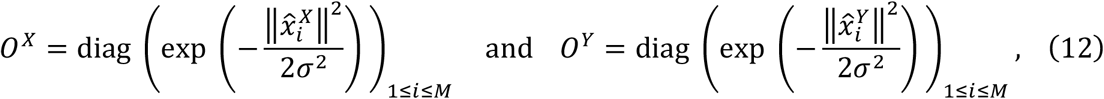

together with the artificial data matrices 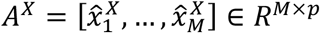 and 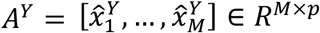, which correspond to the linear terms. Finally, we define:

- 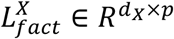 and 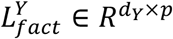, which contain the contributions of each gene (columns) to each source and target latent factor (rows) respectively.
- 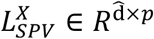 and 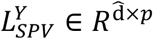 containing the contribution of each gene (columns) to each source and target principal vector (rows) respectively.

These matrices of linear weights can be computed as follows (Supp. Theorem 6.11.):

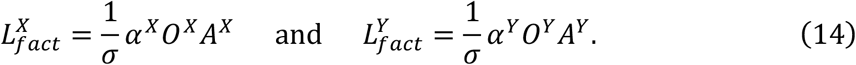

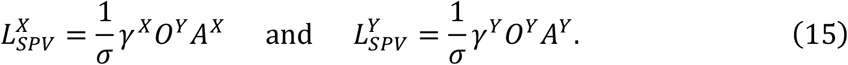

Following a similar protocol, we can compute the contributions of each interaction term to the latent factors and the SPVs, and we refer the reader to Supp. Proposition 6.15 for the complete definition.

To extend this interpretability scheme to other Matérn kernels, e.g., the Laplacian kernels, we proved the orthogonality of linear and interaction terms within the RKHS (Supp. Lemmas 6.16 and 6.17). In order to scale linear and interaction terms, we exploited a heuristic which consists in comparing the amount of non-linearities with results obtained using a Gaussian kernel with same parameter *σ* (Supp. Subsect. 6.5).

### Visualization using Sobolev Principal Vector Interpolation

In order to reduce the SPV pairs to a single vector, we followed the same protocol as in PRECISE and TRANSACT and devised an interpolation scheme (Supp. Material). If we denote by *ϑ_k_* between vectors of the *k^th^* SPV, i.e., 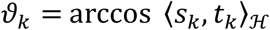, the projection on interpolation time *τ* ∈ [0,1] is defined as

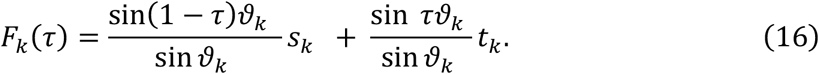

For each SPV, we discretize [0,1] and project the whole data on all the interpolated vectors. Using a Kolmogorov-Smirnov statistics, we compare source and target projected data and return the interpolation time *τ_k_* which minimizes these statistics. We repeat this step for all SPVs and project the data on all the interpolated vectors.

### Gene set enrichment analysis

To connect the linear and interaction weights (Eq. (14) and (15)) to known biological processes, we employ Gene Set Enrichment Analysis (GSEA)^40^. We process the linear and interaction parts in two distinct analyses. First, the linear weights are used as input to a PreRanked analysis with 1000 gene-level permutations. For the interaction terms, we compute enrichment score following a protocol inspired from the original GSEA framework. As interaction terms correspond to pairs of genes, we define for two gene sets *G*_1_ and *G*_2_ the interaction gene set *G*_1_ × *G*_2_ as the set of gene pairs with one gene in *G*_1_ and the other in *G*_2_. To compute the enrichment score of *G*_1_ × *G*_2_, termed *ES*(*G*_1_ × *G*_2_), we rank all interaction terms in decreasing order and assign to each interaction 1 if the interaction is in *G*_1_ × *G*_2_ and −1 otherwise. We then divide the positive values by the number of elements in *G*_1_ × *G*_2_ and the negative values by the number of interactions not in *G*_1_ × *G*_2_. We finally compute the cumulative sum over the ranked set of interactions and define *ES*(*G*_1_ × *G*_2_) as the cumulative sum with the largest absolute value.

Taking inspiration from the original GSEA protocol, we generated the null model by sample permutation: once the scVI model is trained and artificial data is generated, we randomly shuffled the sample labels of latent space values. We then ran the Sobolev Alignment protocol and computed the interaction weights. Although the linear analysis has been carried out with 1 000 permutations, we only performed 100 permutations for the interaction terms, out of computational time considerations. We finally computed the enrichment scores for each permutation and computed the normalized enrichment scores (NES) and FDR values as in the original GSEA.

### Code availability

Sobolev Alignment is available as a Python 3.8 package (<u>GitHub link</u>). Scripts to reproduce the figures from this manuscript are <u>available on GitHub</u>.

## Supporting information

Algorithm derivation

## Authors contribution

S.M.C.M, M.J.T.R., M.L. and L.F.A.W. designed the study. S.M.C.M. performed the experiments. M.J.T.R., M.L. and L.F.A.W. supervised the experiments. S.M.C.M., J.C.S., M.J.T.R and L.F.A.W. analyzed the results. S.M. and M.L. developed the mathematical framework. S.M.C.M. developed the software package. S.M.C.M., J.C.S. and L.F.A.W. analyzed the GSEA results. All authors wrote and approved the manuscripts.

## Acknowledgement

We thank Mirrelijn van Nee (VUmc), Stavros Makrodimitris (Erasmus MC), Chirag Raman (TU Delft), Ahmed Mahfouz (LUMC), Jonas Teuwen (NKI), Bram Thijssen (NKI), Tycho Bismeijer (NKI), Kat Moore (NKI) and Thomas Battaglia (NKI) for useful discussions. We thank Andrea Murachelli (NKI) and Daniel Vis (NKI) for useful technical support. We thank Mark A. van de Wiel (VUmc) for critical reading of the manuscript. We thank the RHPC facility from the Netherlands Cancer Institute (NKI) for the computing infrastructure.

This work was supported by ZonMw TOP grant COMPUTE CANCER [40-00812-98-610 16012].

