## Supplementary Figures for "Identifying commonalities between cell lines and tumors at the single cell level using Sobolev Alignment of deep generative models"

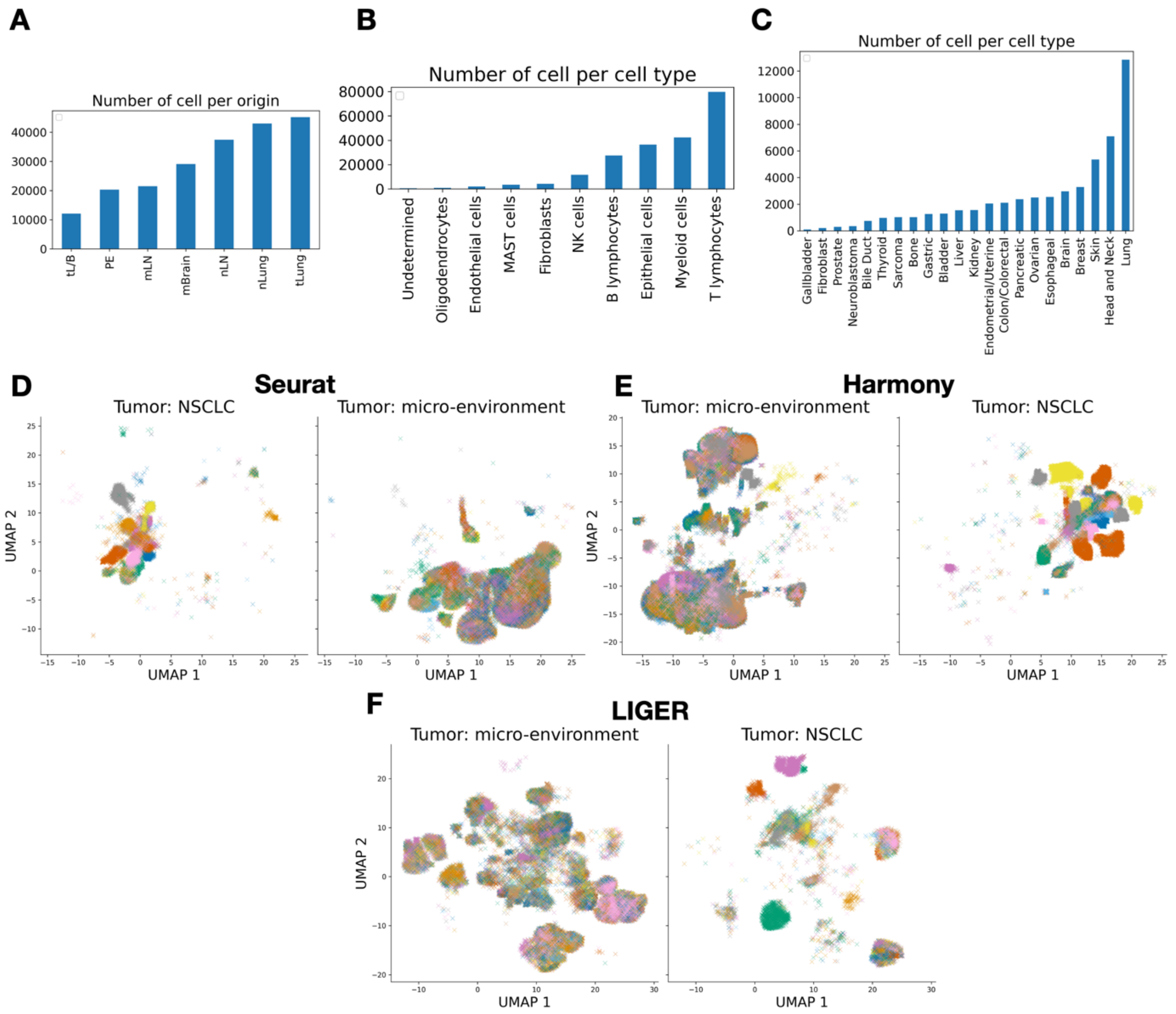


**Extended Figure 1** <u>(Supporting Figure 1) Existence of systematic differences between cell lines and tumors hinders batch effect correction.</u> (**A**) Composition of the *Kim et al*[20] dataset by donors (*L/B and tLung*: primary lung lesion; *PE*: pleural fluids; *mLN*: metastatic lymph node; *mBrain*: brain metastasis; *nLung*: healthy lung; *nLN*: healthy lymph node). (**B**) Composition of the *Kim et al*[20] dataset by cell types. (**C**) Composition of the *Kinker et al*[18] dataset by tissue types. (**D**) UMAP of the Seurat corrected gene expression profiles for patient data (*Kim et al*) divided between epithelial tumor cells (left) and other cells (right), colored by patients. (**E**) UMAP of the Harmony corrected gene expression profiles for patient data colored by patients. (**F**)

UMAP of the LIGER corrected gene expression profiles for patient data colored by patients.

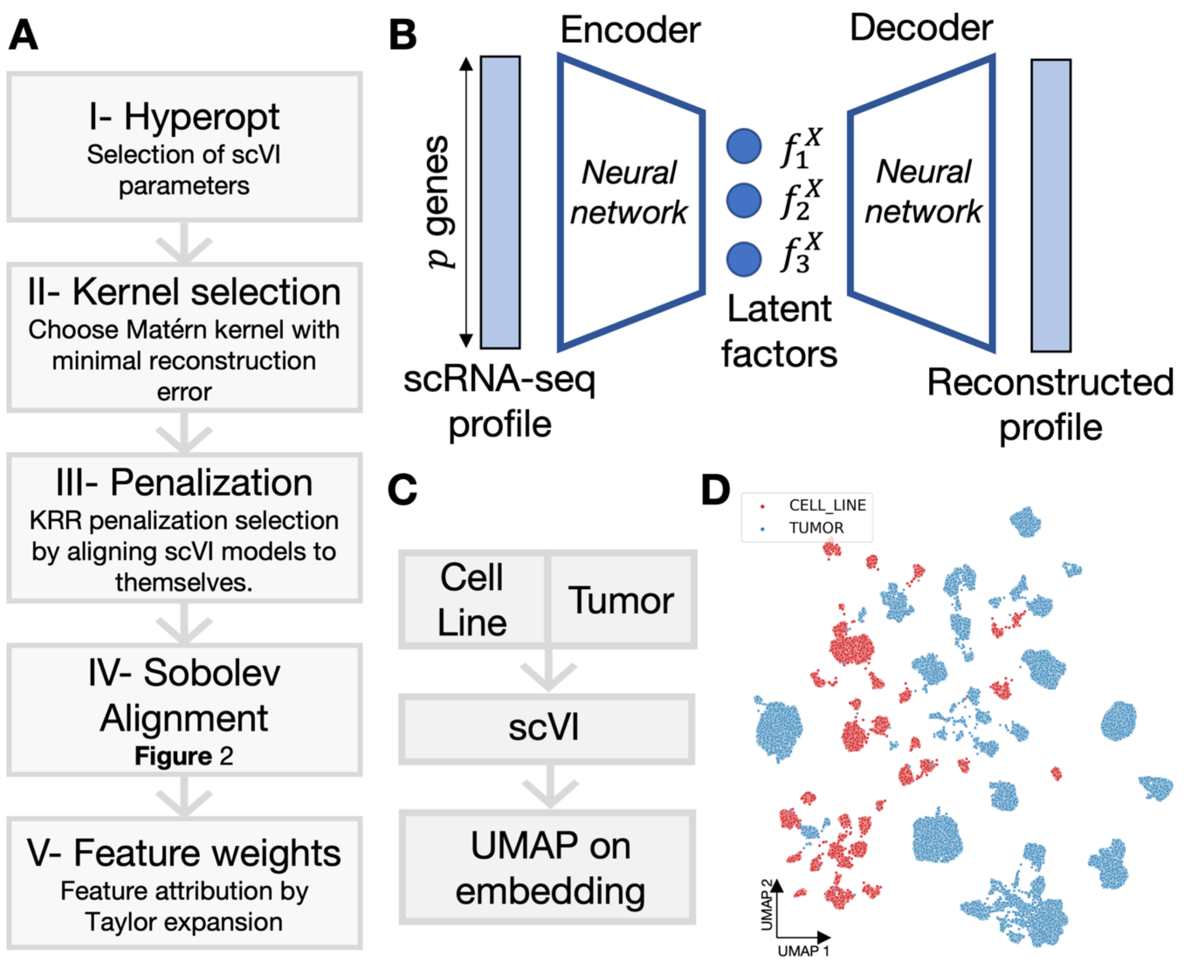


**Extended Figure 2** <u>(Supporting Figure 2) Technical supplement on the Sobolev Alignment algorithm.</u> (**A**) Complete workflow of Sobolev Alignment. In a first step, the hyperparameters of the scVI models (dropout rate, likelihood, weight decay, network architecture, learning rate, learning rate scheduler, early stopping) are set by minimizing the reconstruction error employing Bayesian Optimisation (Hyperopt). In a second step, we set the parameters of the Matérn kernel to be used in Sobolev Alignment. To do so, we first set the scale parameter ($\sigma$) as the median distance observed between source and target single cell profiles. We then train different KRR models on cell line and tumor data with varying values of $\nu$. We use artificial points as training data, and we select $\nu$ as the parameter providing the largest Spearman correlation between the embedding values and the KRR values predicted from the scRNA-seq dataset (not used to train the KRR). In a third step, we set the penalty parameter of the cell line KRR model by aligning the trained cell line scVI model to itself. As we align the exact same model to itself, the SPVs should have a similarity close to one. However, small penalization values would lead to overfitting and therefore decrease this observed similarity – overfitting artifacts have a limited chance to be shared between source and target. We train KRR models with different regularization values and select the lowest value past a certain threshold of self-similarity (by default set to 0.9). We proceed similarly for the tumor regularization parameter. Finally, we perform the whole alignment as explained in **Figure 2**. (**B**) General presentation of an Auto-Encoder. A scRNA-seq profile is used as input into a first neural network, called "encoder", which compresses the data to a small number of "latent factors". These latent factors are then fed into a second neural network, called a "decoder", which maps the latent factors back to the original space. The Auto-Encoder is trained by modifying the weights of both neural networks so that the difference between the input and the output is minimal. A Variational Auto-Encoder (VAE) offers a probabilistic extension of this framework with a more complex architecture which allows the incorporation of prior knowledge; it however still relies on this "encoder-decoder" scheme. (**C**) We trained a single scVI model on the concatenated data (Methods) and used scVI native batch correction to account for batch effect, both within and between datasets. The resulting latent factors were then projected in 2 dimensions using UMAP.

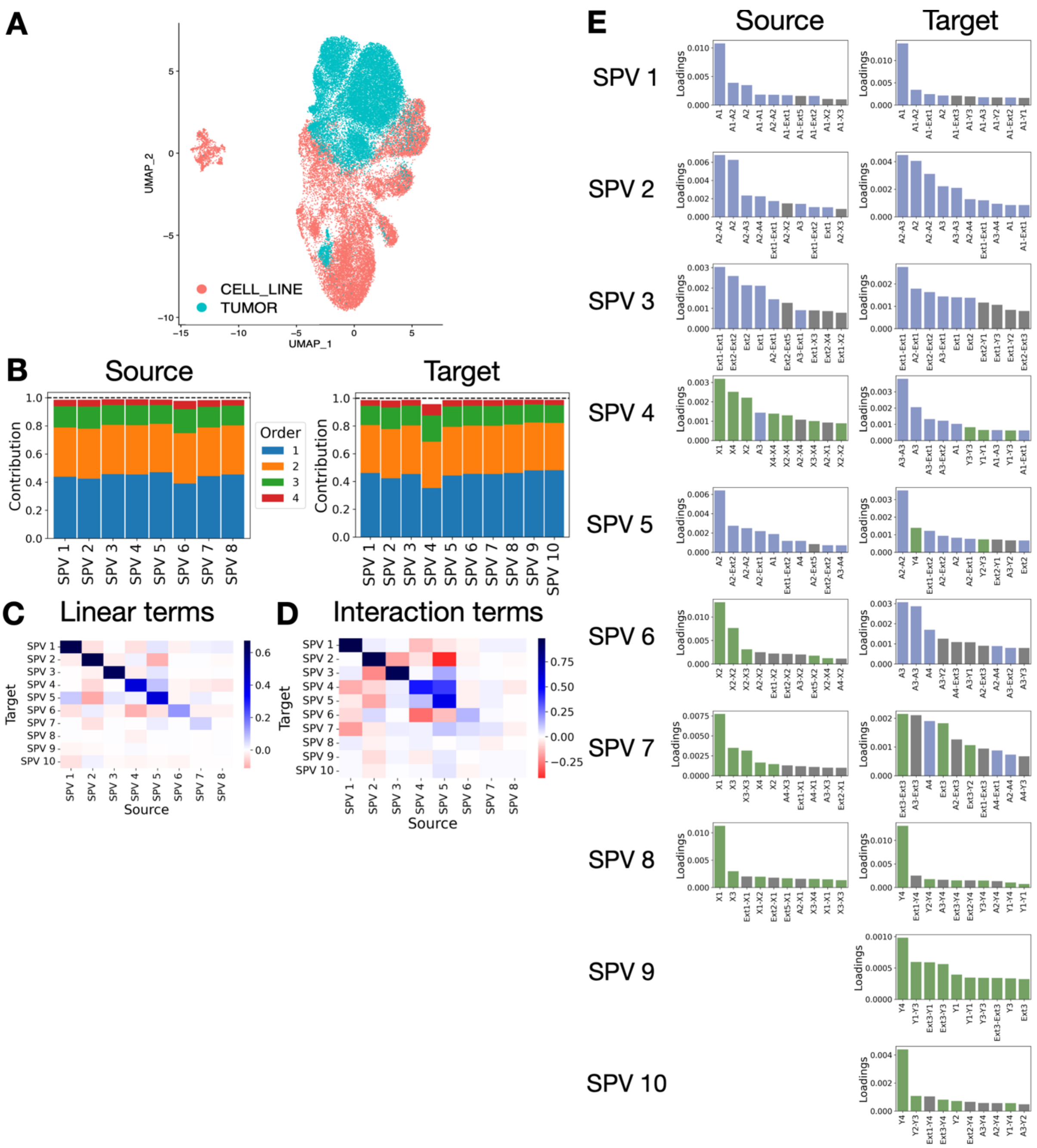


**Extended Figure 3** <u>(Supporting Figure 3) Analysis of model I.</u> (**A**) UMAP obtained after Seurat correction between source (cell lines) and target (tumor). (**B**) Contributions of different order features for source (left) and target (right) SPVs. (**C**) Spearman correlations between the linear weights of source SPVs (x-axis) and target SPVs (y-axis). (**D**) Spearman correlations between the interaction weights of source SPVs (x-axis) and target SPVs (y-axis). (**E**) Square contributions of linear and

interaction features to each SPV, restricted to the top 10 highest contributions and colored as in **Figure 3.**

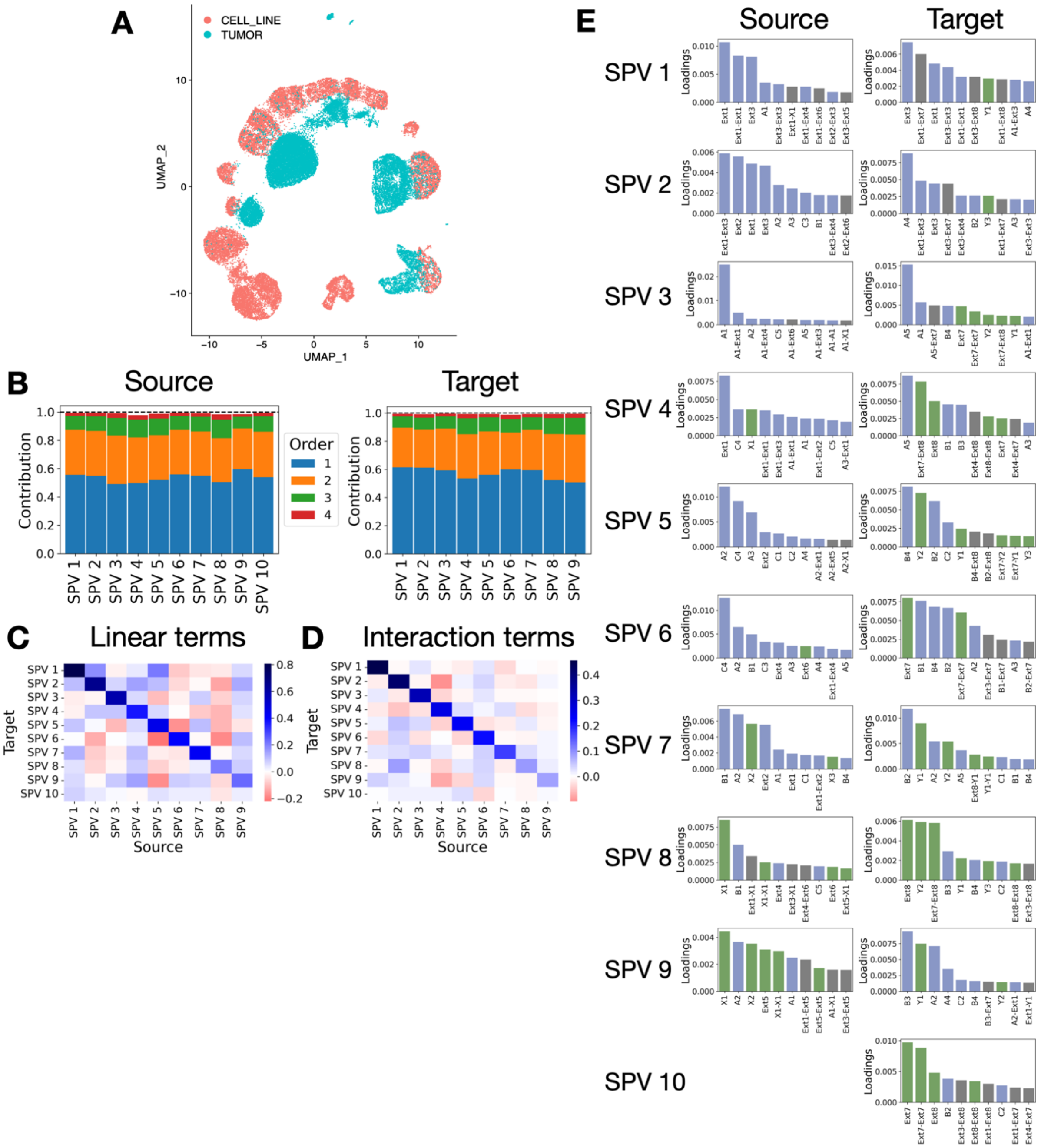


**Extended Figure 4** <u>(Supporting Figure 3) Analysis of model II.</u> (**A**) UMAP obtained after Seurat correction between source (cell lines) and target (tumor). (**B**) Contributions of different order features for source (left) and target (right) SPVs. (**C**) Spearman correlations between the linear weights of source SPVs (x-axis) and target SPVs (y-axis). (**D**) Spearman correlations between the interaction weights of source SPVs (x-axis) and target SPVs (y-axis). (**E**) Square contributions of linear and

interaction features to each SPV, restricted to the top 10 highest contributions and colored as in **Figure 3.**

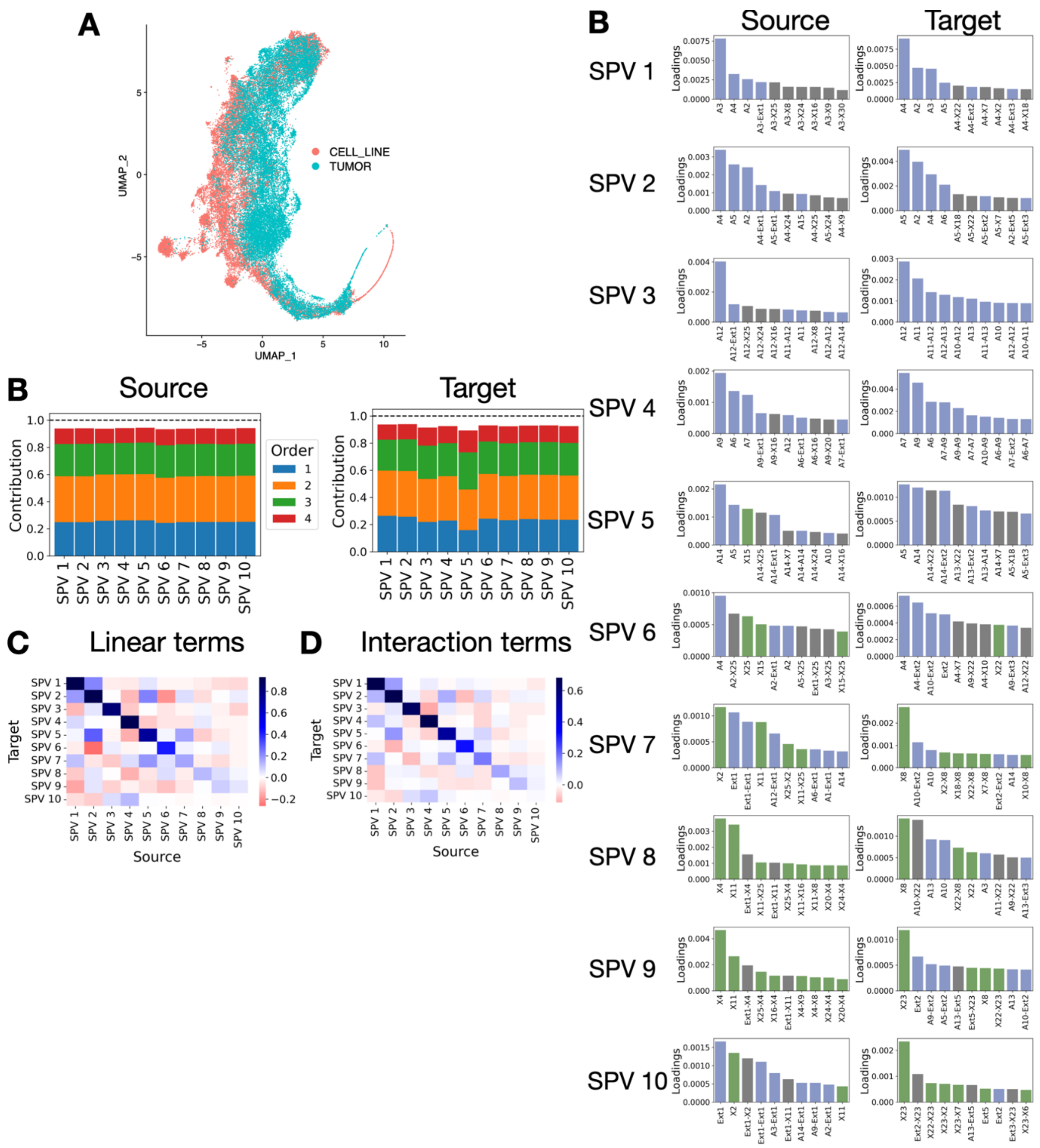


**Extended Figure 5** <u>(Supporting Figure 3) Analysis of model III.</u> (**A**) UMAP obtained after Seurat correction between source (cell lines) and target (tumor). (**B**) Contributions of different order features for source (left) and target (right) SPVs. (**C**) Spearman correlations between the linear weights of source SPVs (x-axis) and target SPVs (y-axis). (**D**) Spearman correlations between the interaction weights of source SPVs (x-axis) and target SPVs (y-axis). (**E**) Square contributions of linear and

interaction features to each SPV, restricted to the top 10 highest contributions and colored as in **Figure 3.**

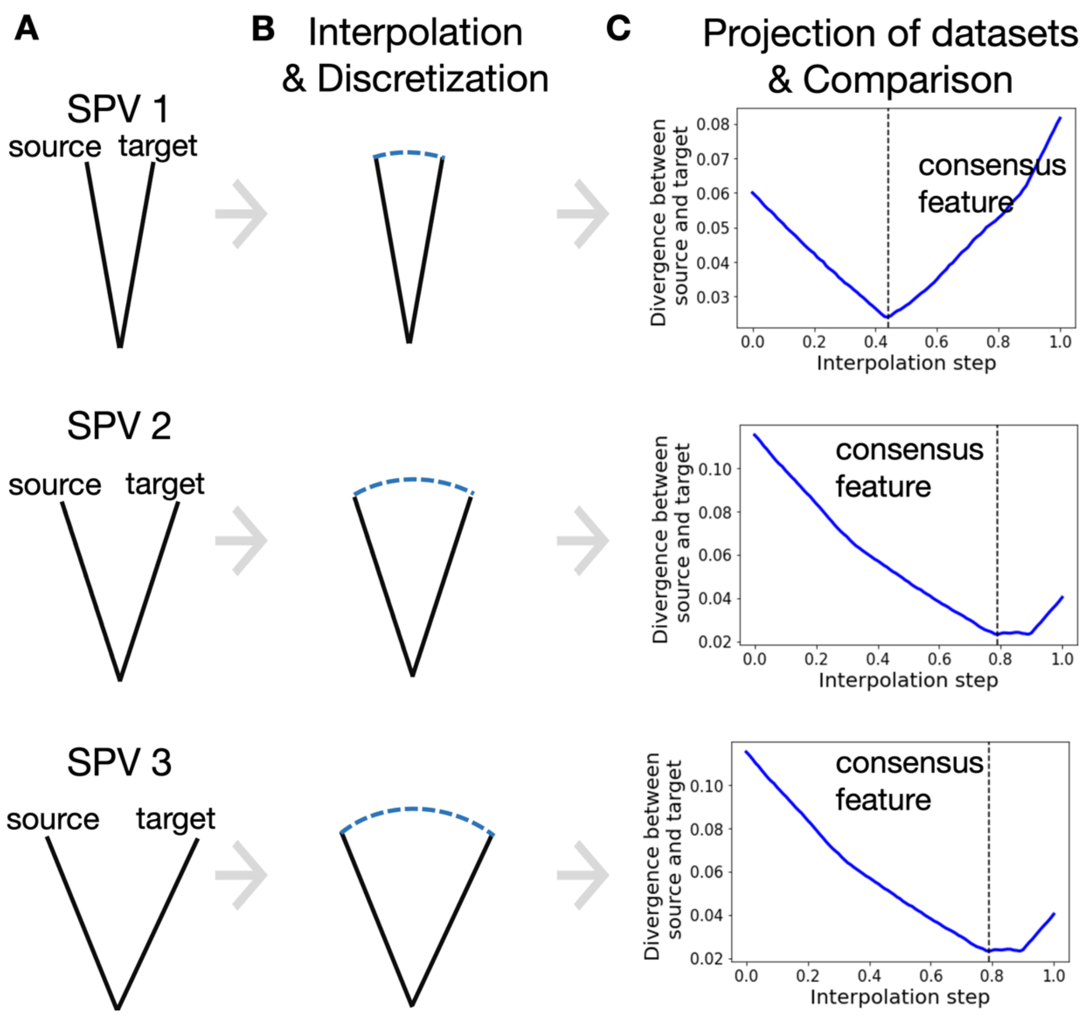


**Extended Figure 6** <u>(Supporting Figure 4) Schematic of the interpolation scheme between similar Sobolev Principal Vectors (SPV).</u> SPVs correspond to pairs of vectors ordered by decreasing similarity. (**A**) When projecting on the top SPVs, it is unclear which of the two vectors to choose: selecting any of the two induces a bias towards either cell lines or tumors. To design a vector which balances the effect of cell lines and tumors, we employ the following interpolation scheme. (**B**) By drawing an arc between the cell line and the tumor vector, we obtain intermediate vectors of same norm. We discretize this arc, e.g., by selecting 100 points equally spaced on this arc, and project cell line and tumor data onto each of these intermediate vectors. (**C**) For each of these intermediate vectors, we compare the cell line and tumor projected data using the Kolmogorov-Smirnoff distance: the lower the distance, the closer the two projected datasets. We select the intermediate vector which minimizes this distance and refer to this vector as the *consensus feature*. This procedure is performed independently for each SPV pair. Such a procedure is inspired from earlier works and stems from an equivalent definition of the geodesic curves in the Grassmann manifold.

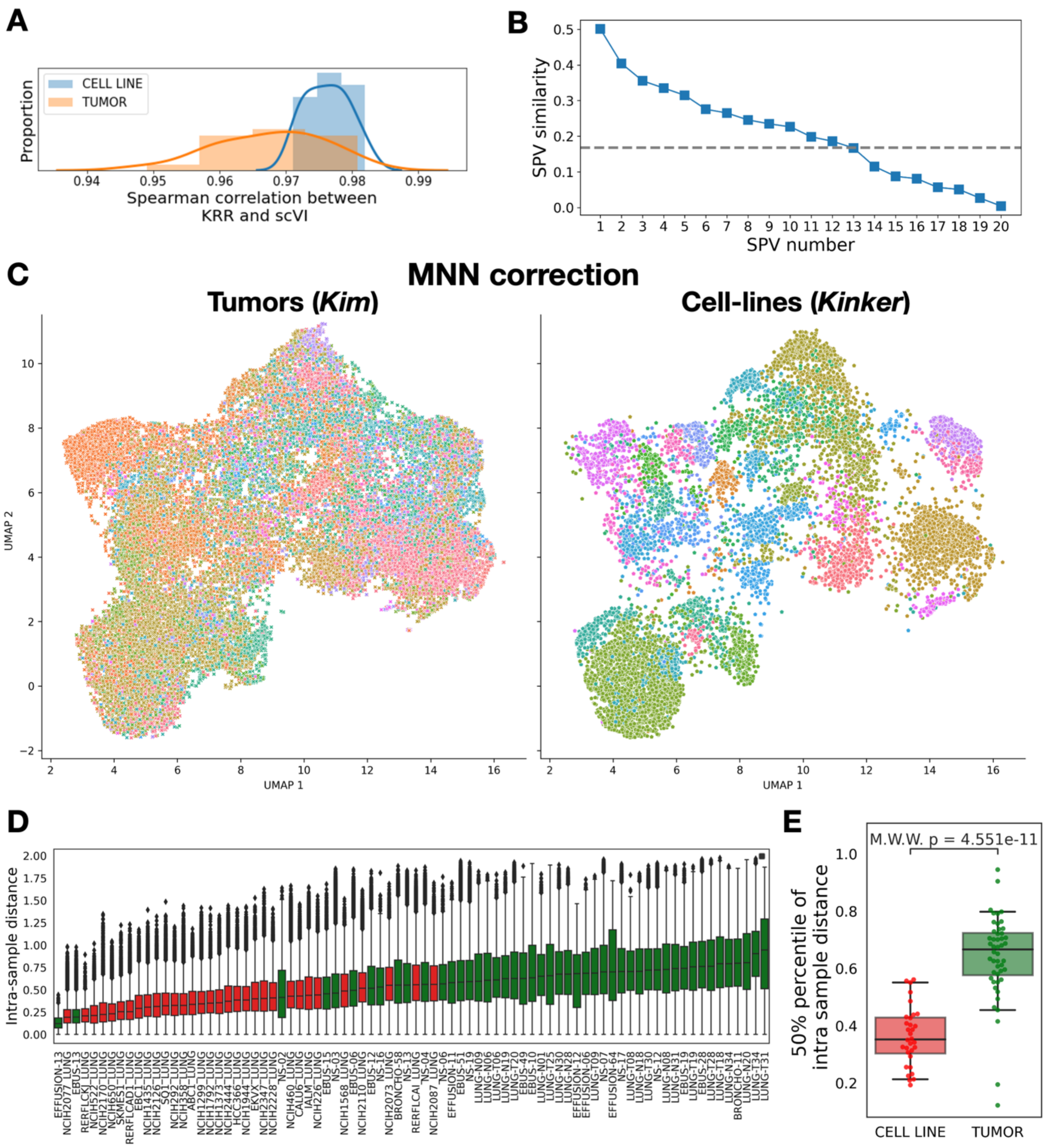


**Extended Figure 7** <u>(Supporting Figure 4) Sobolev Alignment allows an effective co-clustering of cell lines and tumors.</u> (**A**) Histograms of spearman correlations between scVI embeddings and KRR approximations by Falkon. (**B**) Similarity between the Sobolev Principal Vectors (SPV) alongside the maximum similarity value observed after 100 gene permutations (dashed line). (**C**) UMAP visualization of Kinker and Kim datasets after Sobolev Alignment and MNN correction (**Figure 4**C) colored by patients (left) and cell line (right). (**D**) Boxplots of distances between cells from the same patient or cell line. Distances are computed as cosine distances between the cells embedded using Sobolev Alignment and MNN correction. (**E**) Median intra-sample distances observed in panel E.

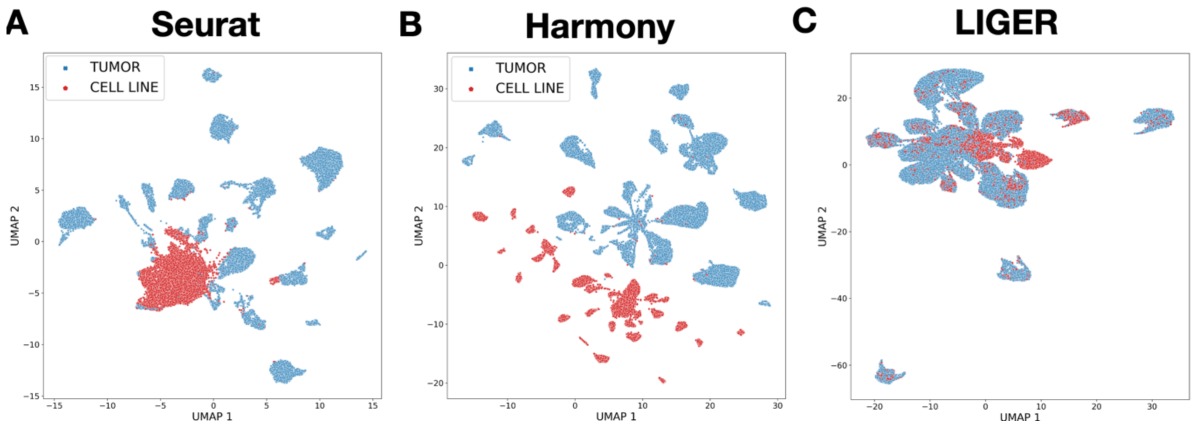


**Extended Figure 8** <u>(Supporting Figure 4) Comparison of integration obtained using standard batch-effect correction tools.</u> We performed the same integration task using 3 state-of-the-art batch effect correction tools. After integration with each method, we projected the data in two dimensions using UMAP. (**A**) Results obtained using Seurat v3. (**B**) Results obtained using Harmony. (**C**) Results obtained using LIGER.

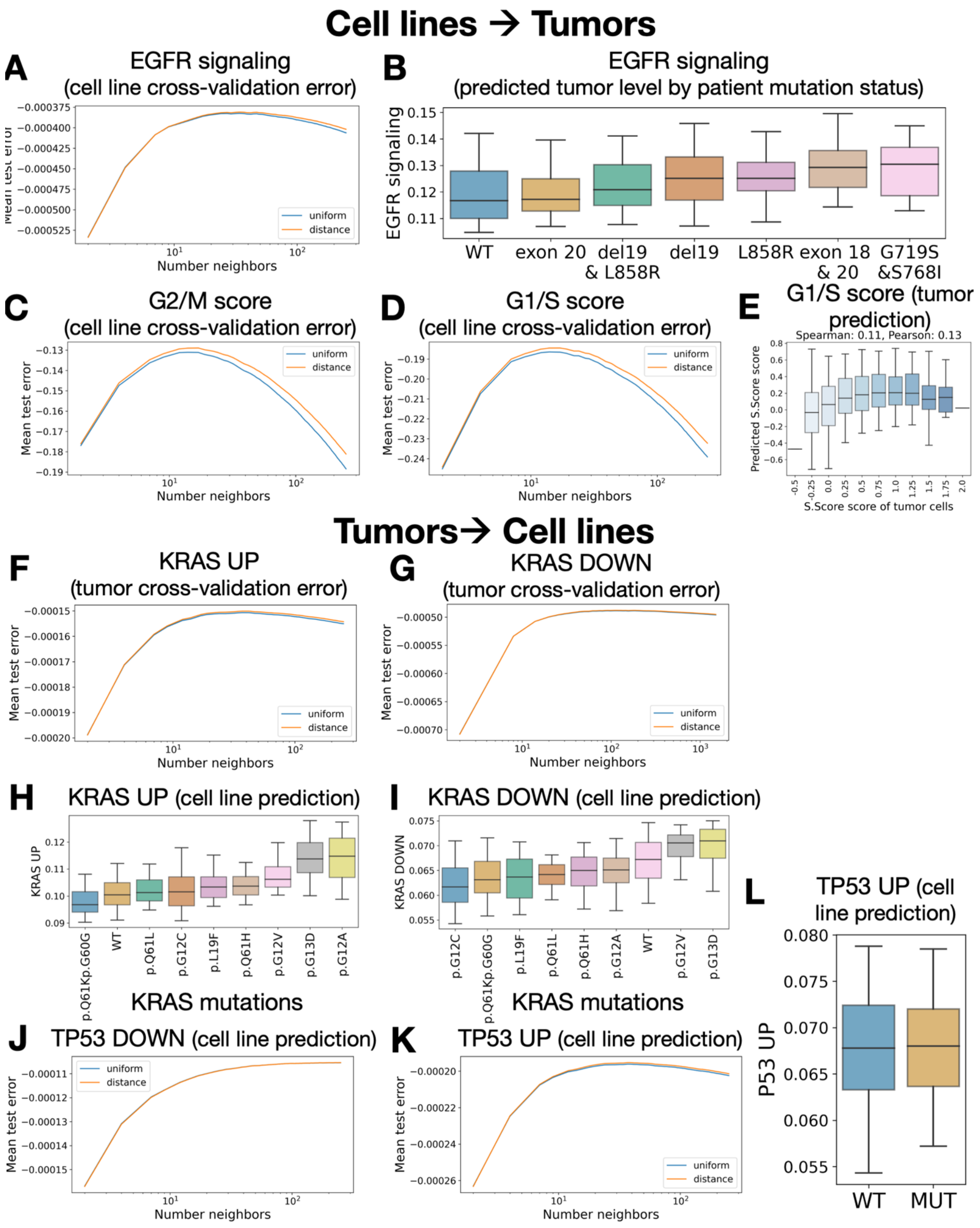


**Extended Figure 9** <u>(Supporting Figure 4) External biomarkers validation of Sobolev Alignment.</u> (**A**) 10-fold cross-validation negative mean squared error (NMSE) obtained on cell line data when training k-Nearest-Neighbors (kNN) regression models on "EGFR signaling" levels (Reactome) for various numbers of neighbors. "Uniform"

indicates that neighbors were similarly weighted for prediction, while “distance” indicates an inverse-distance weighting. (**B**) Predicted EGFR signaling level for tumor cells broken down by EGFR mutations. (**D**) 10-fold cross-validation NMSE on cell lines when training kNN regression models on G2/M scores. (**E**) 10-fold cross-validation NMSE on cell lines when training kNN regression models on S-phase scores. (**F**) Boxplots of predicted S-phase score on tumor cells compared to S-phase scored measured using Seurat v3 cell-cycle regression tool. Spearman and Pearson correlation are computed between the continuous (non-binned) values. (**G**) 10-fold cross-validation NMSE on tumors when training kNN regression models on “KRAS UP” levels (Hallmarks). (**H**) Predicted “KRAS UP” level for tumor cells broken down by KRAS mutations.

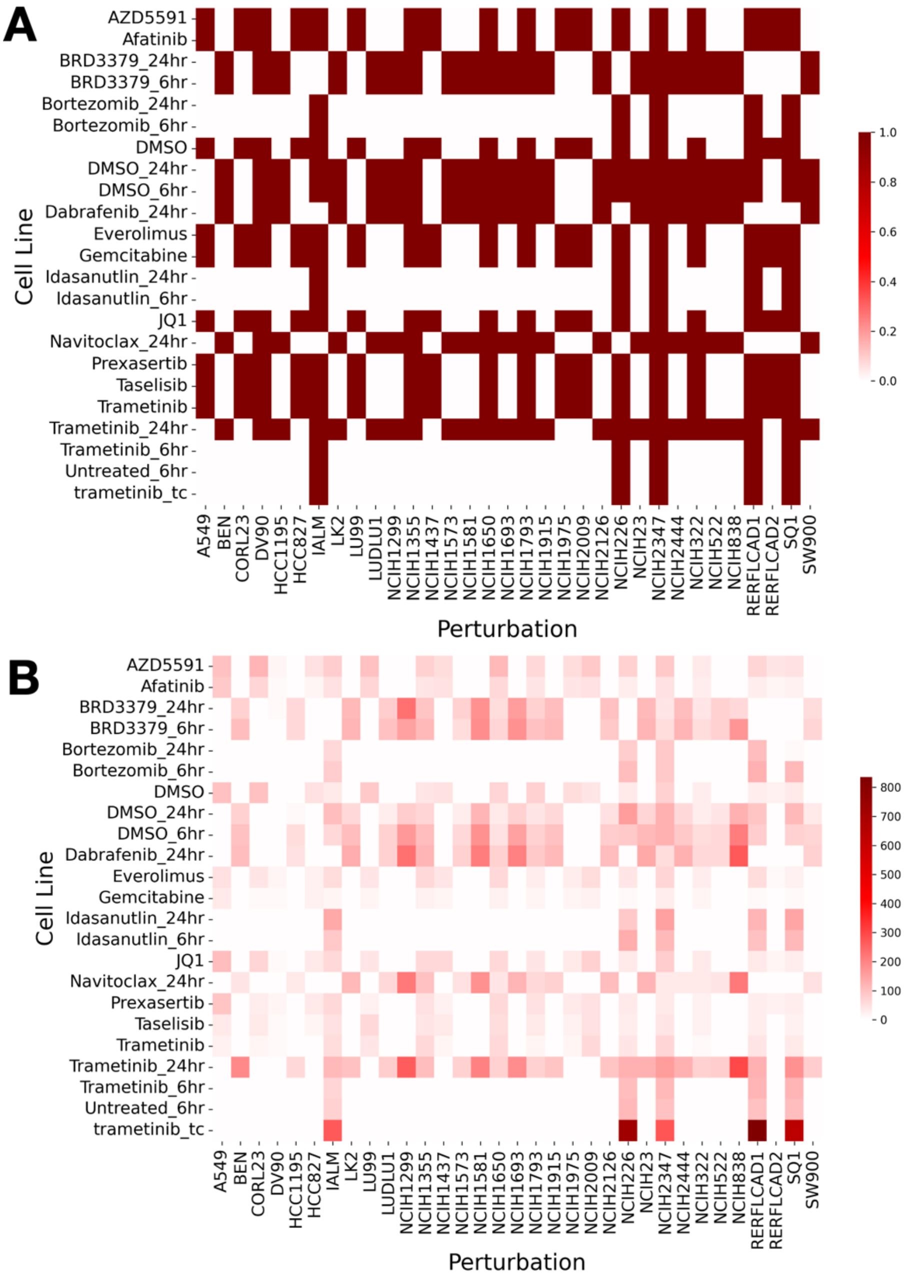


**Extended Figure 10** <u>(Supporting Figure 6) Structure of the employed perturbation screen (McFarland dataset).</u> (**A**) Heatmap indicating whether a cell line (column) has been screened for a certain anti-cancer compound (row). (**B**) Heatmap indicating the number of cells retrieved for each condition.

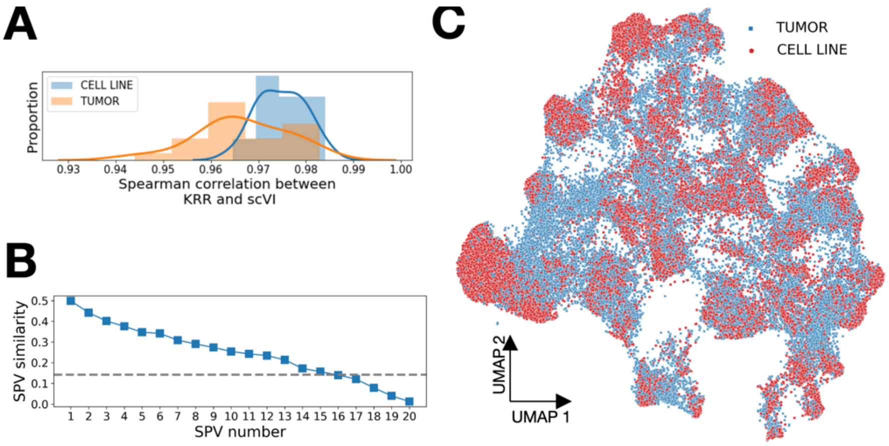


**Extended Figure 11** <u>(Supporting Figure 6) Sobolev Alignment between the McFarland and Kim datasets.</u> (**A**) Histograms of spearman correlations between scVI embeddings and KRR approximations by Falkon. (**B**) Similarity between the Sobolev Principal Vectors (SPVs) alongside the maximum similarity value observed after 100 gene permutations (dashed line). (**C**) UMAP of cell lines and tumors after projection on the top SPVs, interpolation and MNN correction.

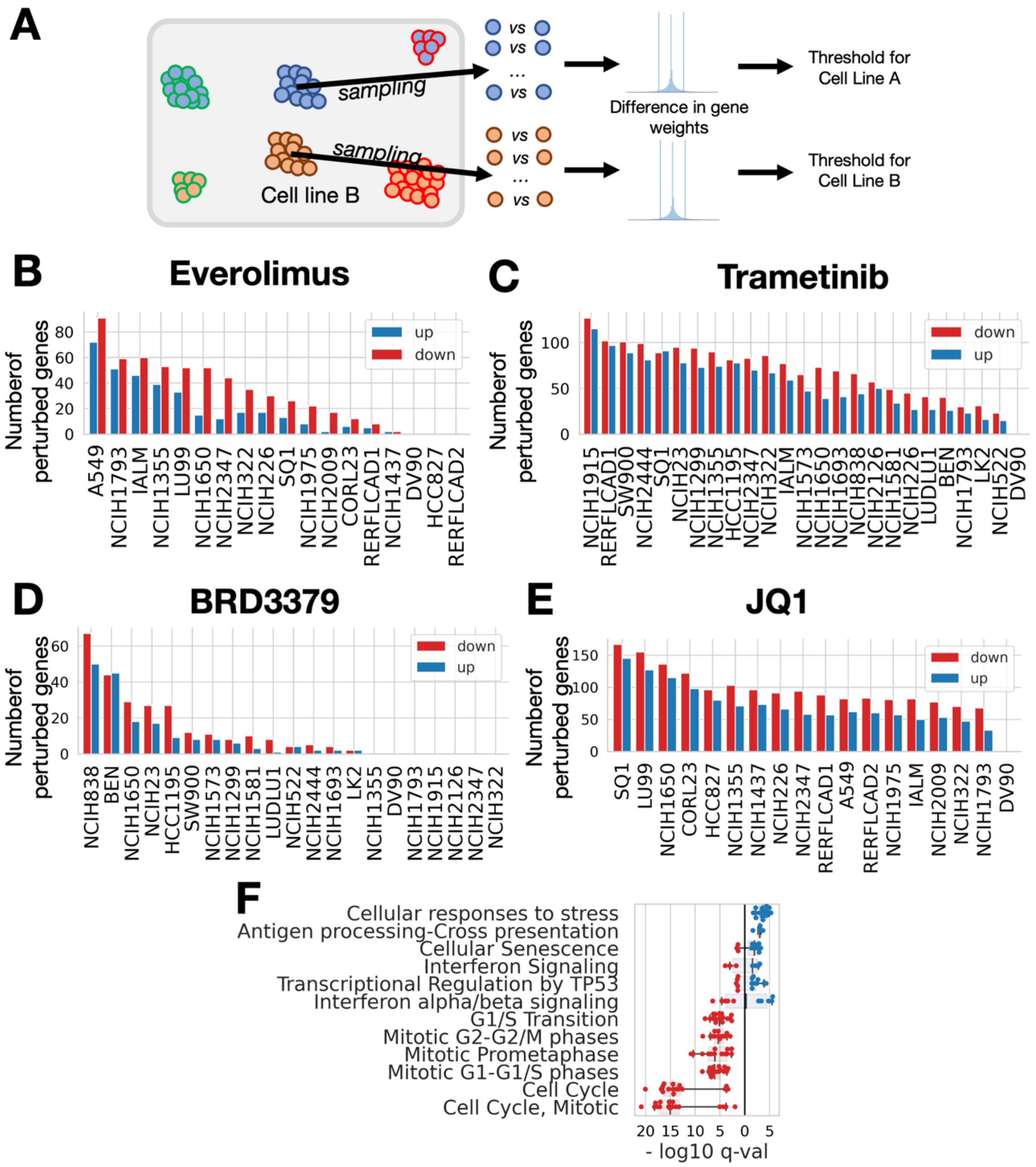


**Extended Figure 12** <u>(Supporting Figure 6) Analysis of drug perturbations after projection on SPVs common between cell lines and tumors.</u> (**A**) To determine the significant effect size threshold, we used the DMSO-treated cells. For each cell line, we randomly sampled 1000 pairs of cells from the pool of DMSO-treated cells and computed the absolute difference in gene weights for all genes. We then took the 95% percentile of all these differences as our effect-size threshold. A gene up- or down-regulation is therefore deemed significant if the Mann-Whitney FDR-corrected p-value is below 0.05 and if the effect size lies above random sampling in DMSO-treated cells. (**B**) Number of perturbed genes, for each cell line, after Everolimus induction. (**C**) Number of perturbed genes for each cell line after Trametinib induction. (**D**) Number of perturbed genes for each cell line after BRD3379 induction. (**E**) Number of perturbed genes for each cell line after JQ1 induction (BRD4-inhibitor). (**F**) Boxplot of q-values obtained when analyzing JQ1 using the Reactome gene sets.
