## Supplementary material for "Identifying commonalities between cell lines and tumors at the single cell level using Sobolev Alignment of deep generative models": Algorithm derivation

*Mathematical derivation and technical details*

Soufiane M.C. Mourragui[1,2], Joseph C. Siefert[1,3], Marcel J.T. Reinders[2,4], Marco Loog[2,5,6], and Lodewyk F.A. Wessels[1,2,6,7]

[1]Computational Cancer Biology, Division of Molecular Carcinogenesis, Oncode Institute, the Netherlands Cancer Institute, Amsterdam, The Netherlands.
[2]Faculty of EEMCS, Delft University of Technology, The Netherlands.
[3]Division of Oncogenomics, the Netherlands Cancer Institute, Amsterdam, The Netherlands
[4]Leiden Computational Biology Center, Leiden University Medical Center, Leiden, The Netherlands.
[5]Department of Computer Science, University of Copenhagen, Copenhagen, Denmark.
[6]Shared last authors

In this supplementary note, we present our approach, Sobolev Alignment, into greater technical and mathematical details. In particular, we provide a full mathematical derivation for the different computational steps presented in the main text. Technical details to replicate our results are also presented here. The reader will find the following elements:

- An introduction to kernel methods and the different mathematical ingredients useful to comprehend our approach are presented in Section 2.
- The different technical choices made for training the different deep generative models are presented in Section 3.
- The complete mathematical derivation is broken down in Sections 4, 5 and 6. The main results of this derivation are highlighted in Theorems 5.2 and 6.11.
- A pseudo-code version of our algorithm is available in Section 7.
- A summary of notations (glossary) is available in Section 8.

**7 Algorithm 15**

**8 Glossary 15**

### 1 Notations

In all of this work, we use gene expression profiles characterized by $p$ genes (or features). Elements referring to the source data are characterized by an $X$ subscript/superscript, while elements referring to the target data are characterized by a $Y$ subscript/superscript.
Source data is comprised of $n_X$ samples, yielding a data matrix $\mathcal{X} \in \mathbb{R}^{n_X \times p}$ ; target data is comprised of $n_Y$ samples, yielding a data matrix $\mathcal{Y} \in \mathbb{R}^{n_Y \times p}$.

### 2 Kernel methods and associated feature space

We here briefly review mathematical and technical details on kernel methods useful to understand the workflow and derivation of Sobolev Alignment. For a longer and detailed presentation of these approaches, we refer the interested reader to [1, 2, 3]. In particular, [4] contains references and proofs to all results listed below.

#### 2.1 Kernel and associated feature space (RKHS)

**Definition 2.1** (Positive Definite Kernel). *A kernel $K$ is a function which takes as inputs two samples and return a scalar value, formally: $K : x, y \in \mathbb{R}^p \to \mathbb{R}$.*
*A kernel $K$ is positive-definite (p.d. in short) if, and only if:*

- *$K$ is symmetric, i.e., for all $x, y \in \mathbb{R}^p, K(x, y) = K(y, x)$.*
- *All kernel matrices are positive definite, i.e.,*

$$\forall n \in \mathbb{N}^*, \forall x_1, ..., x_n \in \mathbb{R}^p, \forall \lambda_1, .., \lambda_n \in \mathbb{R}, \quad \begin{cases} \sum_{i=1}^{n} \lambda_i \lambda_j K(x_i, x_j) \geq 0 \\ \sum_{i=1}^{n} \lambda_i \lambda_j K(x_i, x_j) = 0 \implies \lambda_1 = \lambda_2 = .. = 0 \end{cases} . \tag{1}$$

A positive-definite kernel can be implicitly understood as an inner product in an Hilbert space. This property is interesting as it allows to perform linear algebra operations in a higher-dimensional space where direct computation would be potentially intractable. Working in an higher-dimensional space allows to incorporate non-linearities which is often needed to model complex processes. This property is usually referred to as the "kernel-trick", and is formalized by the *Aronszajn theorem.*

**Theorem 2.2** (Aronszajn). *$K$ is positive-definite if and only if there exists a Hilbert space $\mathcal{H}$ and a mapping function $\phi : \mathbb{R}^p \mapsto \mathcal{H}$ such that:*

$$\forall x, y \in \mathbb{R}^p, \quad K(x, y) = \langle \phi(x), \phi(y) \rangle_{\mathcal{H}}. \tag{2}$$

This feature space is a **Reproducing Kernel Hilbert Space**, or RKHS in short, and is an intrinsic property of the kernel [1]. By construction, an RKHS corresponds to a functional space and each sample embedding $\phi(x)$ (for $x \in \mathbb{R}^p$) can be understood as a function $\mathbb{R}^p \to \mathbb{R}$. The elements of $\mathcal{H}$ are both vectors in a high-dimensional space and functions. This duality allows us to approximates encoder functions in such an RKHS, and also apply linear algebra and compare encoders as vectors. This is formalized by the **Reproducing property**.

**Proposition 2.3** (Reproducing property). *Let $K$ be a p.d. kernel with RKHS $\mathcal{H}$. Then $\mathcal{H}$ is a set of functions, i.e. $\mathcal{H} \subset \{f : \mathbb{R}^p \to \mathbb{R}\}$ with the two following properties:*

[1] The RKHS is actually defined by adding two more hypothesis, which we do not list for sake of simplicity. We refer the reader to [2] for a complete construction of the RKHS.

- $\forall x \in \mathcal{H}, \quad K_x \,\hat{=}\, \quad y \mapsto K(y,x) \quad \in \mathcal{H}$
- $\forall f \in \mathcal{H}, \forall x \in \mathbb{R}^p, \quad f(x) \;=\; \langle K_x, f \rangle_{\mathcal{H}}$

We will now define the three (p.d.) kernels we employ in our study, alongside their feature spaces.

#### 2.2 Gaussian, Matérn and Laplacian kernel and associated RKHS

We consider three different kernels in our work. We here define them and we will show in a subsequent part how these three are related.

**Definition 2.4** (Laplacian kernel)**.** *Let $\sigma > 0$ we define the Laplacian kernel $K_\sigma^L$ on $\mathbb{R}^p$ as:*

$$\forall x, y \in \mathbb{R}^p, \quad K_\sigma^L(x,y) \;=\; \exp\left(-\frac{\|x-y\|}{\sigma}\right). \tag{3}$$

**Definition 2.5** (Matérn kernel [5])**.** *Let $\nu > 0$ and $\sigma > 0$, we define the Matérn kernel $K_{\nu,\alpha}^M$ on $\mathbb{R}^p$ as:*

$$\forall x, y \in \mathbb{R}^p, \quad K_{\nu,\sigma}^M(x,y) \;=\; \frac{2^{1-\nu}}{\Gamma(\nu)}\left(\frac{\sqrt{2\nu}\|x-y\|}{\sigma}\right)^{\nu} K_\nu\left(\frac{\sqrt{2\nu}}{\sigma}\|x-y\|\right), \tag{4}$$

*where $\Gamma$ is the Gamma function, and $K_\alpha$ the modified Bessel function of second kind of order $\alpha$.*

A few interesting examples of Matérn kernels are the following:

$$\begin{aligned}
\text{Order 1/2:} \quad k_{\frac{1}{2},\sigma}(x,y) \;&=\; \exp\left(-\frac{\|x-y\|}{\sigma}\right) \\
\text{Order 3/2:} \quad k_{\frac{3}{2},\sigma}(x,y) \;&=\; \left(1+\frac{\sqrt{3}\|x-y\|}{\sigma}\right)\exp\left(-\frac{\sqrt{3}\|x-y\|}{\sigma}\right) \\
\text{Order 5/2:} \quad k_{\frac{5}{2},h}(x,y) \;&=\; \left(1+\frac{\sqrt{5}\|x-y\|}{\sigma}+\frac{5\|x-y\|^2}{3\sigma^2}\right)\exp\left(-\frac{\sqrt{5}\|x-y\|}{\sigma}\right)
\end{aligned} \tag{5}$$

**Definition 2.6** (Gaussian kernel)**.** *Let $\sigma > 0$, we define the Gaussian kernel $K_\sigma^L$ on $\mathbb{R}^p$ as:*

$$\forall x, y \in \mathbb{R}^p, \quad K_\sigma^G(x,y) \;=\; \exp\left(-\frac{\|x-y\|^2}{2\sigma^2}\right). \tag{6}$$

#### 2.3 Equivalence of the three kernels and hyper-parameters

As already hinted at by the first line of Equation (5), the Gaussian, Matérn and Laplacian kernels are related.

**Proposition 2.7** (Equivalence of Gaussian, Matérn and Laplacian kernels)**.** *Let $\sigma > 0$, we have the following equalities:*

$$\textit{Equivalence between Matérn and Laplacian}: \quad K_{\frac{1}{2},\sigma}^M = K_\sigma^L. \tag{7}$$

$$\textit{Equivalence between Matérn and Gaussian}: \quad \lim_{\nu \to +\infty} K_{\nu,\sigma}^M = K_\sigma^G. \tag{8}$$

#### 2.4 Relationship between Matérn feature spaces and Sobolev spaces

We first start by a general definition of the RKHS of Matérn kernel, which is related to the Gaussian and Laplacian kernels (Proposition 2.7). Matérn kernels are related to the so-called *Sobolev spaces* which are functional spaces used in various areas of mathematics and physics.

**Definition 2.8** (Weak differentiation operation). *Let $\beta \in \mathbb{N}^p$ and $|\beta| = \sum_{1\leq i\leq n} \beta_i$. Let $f$ be a function from $\mathbb{R}^p$ to $\mathbb{R}$, $\beta$-weakly differentiable. We denote by $D^\beta f$ the $\beta^{th}$weak differential of $f$. In the particular case when $f$ is $|\beta|$-times differentiable, we have the following equality:*

$$D^\beta f \;=\; \frac{\partial^{|\beta|}}{\partial x_1^{\beta_1} ... \partial x_p^{\beta_p}} f. \tag{9}$$

**Definition 2.9** (Space of continuous integrable functions). *We define as $L_2(\mathbb{R}^p)$ the space of continuous functions defined as follows:*

$$L_2(\mathbb{R}^p) \;=\; \left\{ f : \mathbb{R}^p \to \mathbb{R} \mid \int_{\mathbb{R}^p} f(x)^2 dx < +\infty \right\}, \tag{10}$$

*endowed with the following inner-product.*

$$\forall f, g \in L_2(\mathbb{R}^p), \quad \langle f, g \rangle_{L_2(\mathbb{R}^p)} \;=\; \int_{\mathbb{R}^p} f(x) g(x) dx. \tag{11}$$

*It defines a Hilbert space[2], which we denote as $L_2$ in the sequel. .*

Using these two bricks, we define a Sobolev spaces as follows:

**Definition 2.10** (Sobolev spaces). *Let $s > 0$ be an integer. We define the Sobolev space of order $s$, denoted noted $W_2^s$ as:*

$$W_2^s \;=\; \left\{ f \in L_2 \quad \mid \sum_{\beta\in\mathbb{N}^p \mid |\beta|\leq s} \left\| D^\beta f \right\|_{L_2}^2 < +\infty \right\}, \tag{12}$$

*endowed with the following inner product:*

$$\forall f, g \in W_2^s, \quad \langle f, g \rangle_{W_2^s} \;=\; \sum_{\beta\in\mathbb{N}^p \mid |\beta|\leq s} \left\langle D^\beta f, D^\beta g \right\rangle_{L_2}. \tag{13}$$

These Sobolev spaces can approximate any function in $L_2$, as shown by the following proposition.

**Proposition 2.11** (Density of $W_2^s$ in $L_2$). *Let $f \in L_2$ and $s > 0$ be an integer. There exists $f_1, f_2, ... \in W_2^s$ such that:*

$$\| f_n - f \|_{L_2} \xrightarrow[n\to+\infty]{} 0 \tag{14}$$

**Proposition 2.12** (Matérn feature space). *Let $\nu > 0$ and $\sigma > 0$. If $\nu + \frac{p}{2}$ is an integer, then $K_{\nu,\sigma}^M$ has for RKHS $\mathcal{H}_{\nu,\sigma} = W_2^{\nu+\frac{p}{2}}$, and the associated norms are equivalent, i.e. there exists $c_1, c_2 > 0$ such that:*

$$\forall f \in W_2^s, \quad c_1 \|f\|_{W_2^s} \;\leq\; \|f\|_{\mathcal{H}_{\nu,\sigma}} \;\leq\; c_2 \|f\|_{W_2^s}. \tag{15}$$

[2] $L_2$ is actually a quotient space defined up to an equivalent class related to the Lebesgue-measure used on $\mathbb{R}^p$. We defined it here as a functional space for sake of clarity.

The result of Proposition 2.12 shows that the feature space associated to the Matérn kernel is equivalent to a Sobolev space. Any kernel-based algorithm which employs the Matérn kernel therefore implicitly operates on functions which lie on a Sobolev space.
By definition, the Sobolev inner product of order $s$ between two functions compares all the derivatives of order $s$ or lower using an Euclidean distance. Due to this natural (and standard) way of comparing functions, coupled with the density argument from in $L_2$ (Proposition 2.11), we decided to use Sobolev spaces to approximate our encoder functions, and therefore turned to Matérn kernel machines. Although directly using the $L_2$ space would have been ideal, we were here hindered by the fact that $L_2$ is **not** an RKHS and therefore not amenable to approximation by kernel machines.

### 3 Training of Variational Auto-Encoders (VAE)

**Definition 3.1** (Space of probability measures)**.** *We define by $P(\mathbb{R}^p)$ the space of all Borel probability measures on $\mathbb{R}^p$.*

**Definition 3.2** (VAE models)**.** *We define the following two sets of models:*

$$\begin{array}{ll} \mu^X : \mathbb{R}^p \mapsto \mathbb{R}^{d_X}, \quad \Sigma^X : \mathbb{R}^p \mapsto \mathbb{R}^{d_X} \quad \text{and} \quad g^X : \mathbb{R}^{d_X} \mapsto P(\mathbb{R}^p) \\ \mu^Y : \mathbb{R}^p \mapsto \mathbb{R}^{d_Y}, \quad \Sigma^Y : \mathbb{R}^p \mapsto \mathbb{R}^{d_Y} \quad \text{and} \quad g^Y : \mathbb{R}^{d_Y} \mapsto P(\mathbb{R}^p) \end{array} . \tag{16}$$

*$g^X$ and $g^Y$ outputs probability distributions on $\mathbb{R}^p$ and are referred to as decoders (or stochastic decoders). The families $(\mu^X, \Sigma^X)$ and $(\mu^X, \Sigma^X)$ are referred to as encoders (or stochastic encoders).*

The two functions $g^X$ and $g^Y$ can be tailored to specific problems. In the particular case of scVI, the decoder is constructed based on a zero-inflated negative binomial (ZINB), which models well the scRNA-seq data structure.

**Definition 3.3** (Latent factors sampling)**.** *We assume that the latent factors follow a multivariate normal prior, i.e.,*

$$z_X \sim \mathcal{N}(0_{d_X}, I_{d_X}) \quad \text{and} \quad z_Y \sim \mathcal{N}(0_{d_Y}, I_{d_Y}). \tag{17}$$

*Given two datapoints $x \in \mathbb{R}^p$ and $y \in \mathbb{R}^p$, its latent factors are computed by sampling from the following multivariate normal posterior:*

$$z_X|x \sim \mathcal{N}\left(\mu^X(x), \operatorname{diag}\left(\Sigma^X(x)\right)\right) \quad \text{and} \quad z_Y|y \sim \mathcal{N}\left(\mu^Y(y), \operatorname{diag}\left(\Sigma^Y(y)\right)\right). \tag{18}$$

Considering a multivariate normal prior over the latent factors is a standard design choice, particularly attractive due to the so-called re-parametrization trick [6]. The original paper of scVI ([7]) contains additional information on the specific model we used.

### 4 Comparing encoders by Kernel Ridge Regression approximation

The VAE models presented in Section 3 concentrate a high-dimensional signal into a few latent factors. Understanding how the latent factors from the source model compare to the ones from the target model is however a difficult task. As presented in the main text, we propose to approximate each VAE model by means of Matérn kernel machines and present here our approach in greater mathematical details.
Our approach stems from the rationale that a wide class of function can be approximated by a Matérn kernel machine, provided enough data-points are available – we say that the Matérn kernel is a universal approximator. As generative models can generate their own data points,

this allows us to exploit the consistency of kernel machines to our advantage by attempting to get as close as possible to the asymptotic limit.
Once the functions have been approximated, we can work in the Matérn kernel feature space (i.e. Sobolev space, Proposition 2.12) and align these functions using the Representer Theorem formulation of the approximated functions.

**Definition 4.1** (Mean function of the VAE)**.** *As explicited in Definition 3.2, the encoder of a VAE is parametrized by two sets of functions: one for the means and one for the standard-deviations. We here consider the mean embedding given to each sample and define, following notations from Equation(16):*

$$\forall t \in \{X, Y\}, \ \forall i \in \{1, .., d_t\}, \quad f_i^t = \mu_i^t. \tag{19}$$

Following the scVI model, we here assume that the latent factors follow a multivariate-normal prior.
We have two sets of encoders we want to align – we will approximate them using two distinct Matérn kernel ridge regression defined as follows.

#### 4.1 Definition of Kernel Ridge Regression

In this subsection, we succintly present some results about Kernel Ridge Regression. For the sake of presentation only, we refer to an hypothetical dataset $\mathcal{D} = \{(\widehat{x}_1, \widehat{z}_1), .., (\widehat{x}_N, \widehat{z}_N)\}$ with $\widehat{x}_i \in \mathbb{R}^p$ and $\widehat{z}_i \in \mathbb{R}$. This dataset does not refer to cell lines or tumors and is given purely for illustrative purposes.

**Definition 4.2** (Matérn Kernel Ridge Regression (KRR))**.** *We approximate a function $f : \mathbb{R}^p \to \mathbb{R}$ by performing Kernel Ridge Regression with the Matérn kernel $K^M_{\nu,\sigma}$. This yields, for $\lambda > 0$, the function $\theta^* \in \mathcal{H}_{\alpha,h}$ solution of:*

$$\theta^* \quad = \quad \underset{\theta \in \mathcal{H}_{\alpha,h}}{\operatorname{argmin}} \ \frac{1}{N} \sum_{i=1}^{N} \left(f\left(\widehat{x}_i\right) - \theta\left(\widehat{x}_i\right)\right)^2 \ + \ \lambda \|\theta\|^2_{\mathcal{H}_{\nu,\sigma}}. \tag{20}$$

The Kernel Ridge Regression problem from Definition 4.2 can be solved in closed form.

**Proposition 4.3** (Solution of KRR)**.** *The solution of Equation (20) is:*

$$\theta^* \ = \ \sum_{k=1}^{N} \alpha_k K^M_{\nu,\sigma}\left(\widehat{x_k}, \cdot\right), \quad \textit{with} \begin{cases} K \ = \ \left(K^M_{\nu,\sigma}\left(\widehat{x}_i, \widehat{x}_j\right)\right)_{1 \le i,j \le N} \\ \widehat{z} \ = \ [\widehat{z_1}, .., \widehat{z_N}] \\ \alpha \ = \ \left(K + \lambda N I_N\right)^{-1} \widehat{z} \end{cases} \tag{21}$$

Solving the problem from Definition (4.2) therefore requires inverting a $N \times N$ matrix, which becomes intractable as soon as $N$ reaches approximately a few tens of thousands. To scale our approach to millions of samples, we exploit recent advances in kernel machines such as Falkon.

**Proposition 4.4** (Falkon approximation of KRR)**.** *The solution of Equation (20) can be approximated by:*

$$\hat{\theta} \ = \ \sum_{k=1}^{M} \alpha_k K^M_{\nu,\sigma}\left(\widehat{x}_k, \cdot\right), \tag{22}$$

*with $M < N$ and $\alpha$ coefficients computed by the Nyström approximation. The $M$ points $\widehat{x}_1, .., \widehat{x}_M$ are referred to as* ***Falkon anchor points****.*

Proposition 4.4 shows that the weight vector $\alpha$ from Proposition 4.3 can be approximated using the Nyström approximation, which is the strategy employed in the Falkon package. We refer the reader to the original Falkon paper for details on how to perform this approximation, as it is not necessary for our derivation. The sum expansion of Equation (22) is the cornerstone of all these methods.
Importantly, the Nyström approximation does **not** correspond to performing KRR by restricting to $M$ subsampled points. The remaining $N - M$ points are influencing the solution and are present in the weights $\alpha$: their influence has been factored in during the training process.

#### 4.2 Approximation of encoders by large-scale KRR

We consider one Matérn kernel $K^M_{\nu,\sigma}$ which we refer to as $K$ in the rest of the text. The RKHS $\mathcal{H}_{\nu,\sigma}$ is also referred to as $\mathcal{H}$ for ease of notation.

**Definition 4.5** (Model Samples)**.** *Let $N \in \mathbb{N}$. Using cell line (resp. tumor) VAE (Section 3), we sample N points, called **Model Samples**, as follows:*

- $z^X_1, .., z^X_N \sim \mathcal{N}(0, I_{d_X})$ *(resp.* $z^Y_1, .., z^Y_N \sim \mathcal{N}(0, I_{d_Y}))$*).*
- *Passing the random points through the decoder, we obtain* $\widehat{x^X_1}, .., \widehat{x^X_N} \in \mathbb{R}^p$ *(resp.* $\widehat{x}^Y_1, .., \widehat{x}^Y_N$*).*
- *We pass these sample points into the encoders to get the embeddings* $\widehat{z^X_1}, .., \widehat{z}^X_N \in \mathbb{R}^{d_X}$ *(resp.* $\widehat{z}^Y_1, .., \widehat{z}^Y_N \in \mathbb{R}^{d_Y}$*).*

A VAE is not bijective: we should therefore expect that the $\widehat{z}^X_i$ will differ from the $z^X_i$. Since we are here interested to approximate the encoder functions, we computed the output of the Model Points by each encoder function. Furthermore, using the latent values sampled from the multivariate normal prior distribution would have led to errors in the approximation due to the stochasticity of the VAE: the random points are sampled from a distribution specific to each latent value and are not deterministic.

**Definition 4.6** (KRR Approximation)**.** *We use the Model Samples (Definition 4.5) as training data to train $d_X$ (resp. $d_Y$) Falkon KRR models for cell lines (resp. tumors), which we termed $\theta^X_1, .., \theta^X_{d_X}$ (resp. $\theta^Y_1, .., \theta^Y_{d_Y}$) (Definition 4.2, Propositions 4.3 and 4.4), with $M < N$. We define the two matrices $\alpha^X \in \mathbb{R}^{d_X \times M}$ and $\alpha^Y \in \mathbb{R}^{d_Y \times M}$ as the KRR sample weights:*

$$\forall t \in \{X, Y\}, \ \forall k \in \{1, .., d_t\}, \quad \theta^t_k \ = \ \sum_{i=1}^{M} \alpha^t_{k,i} K\left(\widehat{x}^t_i \ , \ \cdot\right). \tag{23}$$

In Definition 4.6, we assumed here that the $M$ Falkon anchor points correspond to the first $M$ model points. This is always true, up to a permutation.

#### 4.3 Cosine similarity matrix

**Definition 4.7** (Un-normalized cosine similarity)**.** *We define the un-normalized cosine similarity matrices between $X$ and $Y$ as the matrix $\widetilde{\mathbf{M}}_{X,Y}$ :*

$$\widetilde{\mathbf{M}}_{X,Y} \ = \ \left[\langle \theta^X_i, \theta^Y_j \rangle_{\mathcal{H}}\right]_{\substack{1 \le i \le d_X \\ 1 \le j \le d_Y}}, \tag{24}$$

*and the cosine similarity matrices of $X$ (resp. $Y$), denoted $\widetilde{\mathbf{M}}_X$ (resp. $\widetilde{\mathbf{M}}_Y$), as:*

$$\widetilde{\mathbf{M}}_X \ = \ \left[\langle \theta^X_i, \theta^X_j \rangle_{\mathcal{H}}\right]_{\substack{1 \le i \le d_X \\ 1 \le j \le d_X}} \quad \text{and} \quad \widetilde{\mathbf{M}}_Y \ = \ \left[\langle \theta^Y_i, \theta^Y_j \rangle_{\mathcal{H}}\right]_{\substack{1 \le i \le d_Y \\ 1 \le j \le d_Y}}. \tag{25}$$

**Definition 4.8** (Similarity matrices). *We define* $K_X \in \mathbb{R}^{N\times N}$*,* $K_Y \in \mathbb{R}^{N\times N}$ *and* $K_{X,Y} \in \mathbb{R}^{M\times M}$ *as:*

$$\begin{aligned} K_X &= \left(K\left(\widehat{x_i^X}, \widehat{x_j^X}\right)\right)_{1\leq i,j\leq M} \\ K_Y &= \left(K\left(\widehat{x_i^Y}, \widehat{x_j^Y}\right)\right)_{1\leq i,j\leq M} \\ K_{X,Y} &= \left(K\left(\widehat{x_i^X}, \widehat{x_j^Y}\right)\right)_{1\leq i,j\leq M} \end{aligned}. \tag{26}$$

**Proposition 4.9** (Computation of un-normalized cosine similarity matrices). *We have the following equalities:*

$$\begin{aligned} \widetilde{\mathbf{M}}_X &= \alpha^X K_X {\alpha^X}^T \\ \widetilde{\mathbf{M}}_Y &= \alpha^Y K_Y {\alpha^Y}^T \\ \widetilde{\mathbf{M}}_{X,Y} &= \alpha^X K_{X,Y} {\alpha^Y}^T \end{aligned}. \tag{27}$$

*Proof.* We here show the proof for $\widetilde{\mathbf{M}}_X$; the two other equalities follow from the same idea.
We first recall the first reproducing property of the kernel $K$:

$$\forall x, y \in \mathbb{R}^p, \quad \langle K(x,\cdot), K(y,\cdot)\rangle_{\mathcal{H}} = K(x,y). \tag{28}$$

Let $i, j \in \{1,..,d_X\}$, we have:

$$\langle \theta_i^X, \theta_j^X \rangle_{\mathcal{H}} = \sum_{k=1}^{M}\sum_{l=1}^{M} \alpha_{i,k}^X \alpha_{j,l}^X K\left(\widetilde{x}_k^X, \widetilde{x}_l^X\right), \tag{29}$$

using the bi-linearity of the Hilbertian norm, the fact that $\alpha_X$ is real-valued, and the first reproducing property.
Combining Equation (29) with the definition of the similarity matrix $K_X$ (Definition 4.8), we obtain the desired property. ■

Using these notations, we define the **cosine similarity matrix** as follows.

**Definition 4.10** (Cosine similarity matrix). *We define the cosine similarity matrix* $\mathbf{M}$ *as:*

$$\mathbf{M} = \left(\widetilde{\mathbf{M}}_X\right)^{-1/2} \widetilde{\mathbf{M}}_{X,Y} \left(\widetilde{\mathbf{M}}_Y\right)^{-1/2}. \tag{30}$$

### 5 Encoder alignment by kernel Principal Vectors

#### 5.1 General definition of Principal Vectors

We define the **Principal Vectors** (PVs) between source and target VAEs as the pairs of vectors (one from source, one from target) with a maximal inner-product in $\mathcal{H}$.

**Definition 5.1** (Principal Vectors). *Let* $\hat{d} = \min(d_X, d_Y)$*. We define the* $\hat{d}$ *Principal Vectors (PVs)* $(s_1, t_1), .., (s_{\hat{d}}, t_{\hat{d}}) \in \mathcal{H}\times\mathcal{H}$ *as the functions that maximise the similarity between source and target subspaces, i.e.,*

$$\forall k \in \{1,..,\hat{d}\}, \quad s_k, t_k = \underset{\substack{s\in \text{span}\left(\theta_1^X,..,\theta_{d_X}^X\right),\\ t\in\text{span}\left(\theta_1^Y,..,\theta_{d_Y}^Y\right)}}{\text{argmax}} \langle s, t\rangle_{\mathcal{H}} \quad s.t \begin{cases} \langle s,s\rangle_{\mathcal{H}} = \langle t,t\rangle_{\mathcal{H}} = 1 \\ \forall i<k, \ s_i \perp s \\ \forall i<k, \ t_i \perp t \end{cases} \tag{31}$$

#### 5.2 Computation of PVs

**Theorem 5.2** (Matérn PVs)**.** *Let* $\mathbf{M} = U\Sigma V^T$ *be the Singular Value Decomposition (SVD) of the cosine similarity matrix (Definition 4.10). Let's define* $\gamma^X$ *and* $\gamma^Y$ *as:*

$$\gamma^X = U^T \left(\widetilde{\mathbf{M}}_{\mathbf{X}}\right)^{-1/2} \alpha^X \quad \text{and} \quad \gamma^Y = V^T \left(\widetilde{\mathbf{M}}_{\mathbf{Y}}\right)^{-1/2} \alpha^Y \tag{32}$$

*Then the PVs (Definition 5.1) can be computed as follows:*

$$\forall k < \hat{d}, \quad s_k \; = \; \sum_{i=1}^{M} \gamma_{k,i}^X K\left(\hat{x}_i^X, \cdot\right) \quad \text{and} \quad t_k \; = \; \sum_{i=1}^{M} \gamma_{k,i}^Y K\left(\hat{x}_i^Y, \cdot\right). \tag{33}$$

*Proof.* By definition of $s_1, .., s_{\hat{d}}$ and $t_1, .., t_{\hat{d}}$ (Equation (31)), there exist $\xi^X \in \mathbb{R}^{\hat{d} \times d_X}$ and $\xi^Y \in \mathbb{R}^{\hat{d} \times d_Y}$ such that:

$$\forall k \in \{1, .., d\}, \quad s_k \; = \; \sum_{l=1}^{d_X} \xi_{k,l}^Y \theta_k^X \quad \text{and} \quad t_k \; = \; \sum_{l=1}^{d_Y} \xi_{k,l}^Y \theta_k^Y. \tag{34}$$

Using the bi-linearity of the inner product, we have:

$$\forall k, l \in \{1, .., d\}, \quad \begin{cases} \langle s_k, t_k \rangle & = & {\xi_k^X}^T \; \widetilde{\mathbf{M}}_{X,Y} \; \xi_k^Y \\ \langle s_k, s_l \rangle & = & {\xi_k^X}^T \; \widetilde{\mathbf{M}}_X \; \xi_l^X \\ \langle t_k, t_l \rangle & = & {\xi_k^Y}^T \; \widetilde{\mathbf{M}}_Y \; \xi_l^Y \end{cases}. \tag{35}$$

Combining Equations (34) and (35) in Equation (31) yields the following:

$$\forall k \in \{1, .., \hat{d}\}, \quad \xi_k^X, \xi_k^Y \; = \; \underset{\substack{u^X \in \mathbb{R}^{d_X}, \\ u^Y \in \mathbb{R}^{d_Y},}}{\operatorname{argmax}} \quad {u^X}^T \; \widetilde{\mathbf{M}}_{X,Y} \; u^Y \quad \text{s.t} \quad \begin{cases} {u^X}^T \widetilde{\mathbf{M}}_X u^X \; = \; 1 \\ {u^Y}^T \widetilde{\mathbf{M}}_Y u^Y \; = \; 1 \\ \forall i < k, \; {u^X}^T \widetilde{\mathbf{M}}_X \xi_i^X \; = 0 \\ \forall i < k, \; {u^Y}^T \widetilde{\mathbf{M}}_Y \xi_i^Y \; = 0 \end{cases}. \tag{36}$$

$\mathcal{H}$ is an Hilbert space, therefore, assuming that basis functions are linearly independent, $\widetilde{\mathbf{M}}_X$ and $\widetilde{\mathbf{M}}_Y$ are symmetric and positive-definite and thus admit a square-root and an inverse-square-root.
Let's define $\kappa^X = \gamma^X \widetilde{\mathbf{M}}_X^{1/2}$ and $\kappa^Y = \gamma^Y \widetilde{\mathbf{M}}_Y^{1/2}$. The PV definition by linearisation (Equation (36)) is then equivalent to the SVD of $\widetilde{\mathbf{M}}_X^{-1/2} \widetilde{\mathbf{M}}_{X,Y} \widetilde{\mathbf{M}}_Y^{-1/2} = \mathbf{M}$ defined as $\mathbf{M} = U\Sigma V^T$.
We therefore have $\kappa^X = U^T$ and $\kappa^Y = V^T$, which turns to $\xi^X = U^T \widetilde{\mathbf{M}}_X^{-1/2}$ and $\xi^Y = V^T \widetilde{\mathbf{M}}_Y^{-1/2}$.
Using the Representer Theorem (Definition 4.6), we obtained desired formula. ■

**Corollary 5.2.1.** *Using same notation as Theorem 5.2 with* $\mathbf{M} = U\Sigma V^T$*:*

$$\forall k \in \{1, .., \hat{d}\}, \quad \langle s_k, t_k \rangle_{\mathcal{H}} \; = \; \Sigma_{k,k} \in [0, 1]. \tag{37}$$

These values can be understood as the cosine values of angles, called **principal angles**.

### 6 Interpreting the latent factors by Taylor expansion of kernel approximation

We here present our interpretability scheme, which aims at understanding which genes, or combinations thereof, contribute the most to the latent factors. Our scheme relies on the Taylor expansion of the kernel used for the approximation. Unfortunately, to the best of our knowledge, no analytical form of an orthonormal basis for the Matérn RKHS exists in the literature. We are therefore limited to the Gaussian kernel of length-scale $\sigma$, which we refer to as $K_\sigma$ in the sequel ; we refer to its RKHS as $\mathcal{H}_\sigma$.

#### 6.1 Orthonormal basis of Gaussian feature space

We here summarise the construction of the orthonormal basis (ONB) which we exploit. Its complete derivation can be found in [8].

**Definition 6.1** (Univariate basis function). *Let $i \in \{1, .., p\}$ and $k \in \mathbb{N}$. We define the univariate basis function $e_i^k : \mathbb{R}^p \mapsto \mathbb{R}$ as:*

$$\forall x \in \mathbb{R}^p, \quad e_i^k(x) \;=\; \frac{x_i^k}{\sigma^k \sqrt{k!}} \exp\left(-\frac{x_i^2}{2\sigma^2}\right) \tag{38}$$

**Definition 6.2** (Gaussian basis function). *Let $I = (I_1, \ldots, I_p) \in \mathbb{N}^p$, we define the Gaussian basis function $G_I$ as:*

$$\forall x \in \mathbb{R}^p, \quad G_I(x) \;=\; \prod_{i=1}^{p} e_i^{I_i}(x) \;=\; \left[\prod_{i=1}^{p} \frac{x_i^{I_i}}{\sigma^{I_i}\sqrt{I_i!}}\right] \exp\left(-\frac{\|x\|^2}{2\sigma^2}\right). \tag{39}$$

**Proposition 6.3** (Orthonormal basis). *Let $I, J \in \mathbb{N}^p$ and $\mathbb{I}_{I,J}$ be the dirac function (equals to one if all values of $I$ and $J$ are equal, zero otherwise). We have:*

$$\langle G_I, G_J \rangle_{\mathcal{H}_\sigma} = \mathbb{I}_{I,J}, \tag{40}$$

*which shows that the $(G_I)_{I \in \mathbb{N}^p}$ defines an orthonormal family of $\mathcal{H}_\sigma$.*
*This family furthermore defines a basis of $\mathcal{H}$, i.e.,*

$$\forall x, y \in \mathbb{R}^p, \quad K_\sigma(x, y) \;=\; \sum_{I \in \mathbb{N}^p} G_I(x)\, G_I(y) \;=\; \langle \mathcal{G}_\sigma(x)\, \mathcal{G}_\sigma(y) \rangle_{\mathcal{H}_\sigma}, \tag{41}$$

*with $\mathcal{G}_\sigma(x) = \sum_{I \in \mathbb{N}^p} G_I(x)\, G_I$.*

*Proof.* See Theorem 3 and Theorem 4 of *Steinwart et al* [8] for the proof of this proposition. ■

#### 6.2 Feature attribution scores

Using results from Proposition 6.3, we highlight three kinds of Gaussian basis functions (Definition 6.2) of interest: linear, interaction and higher-order interaction terms.

**Definition 6.4** (Dirac vector). *Let $k \in \{1, .., p\}$. We define as $\delta_k \in \mathbb{N}^p$ the vectors of zeros, with a single one at the $k$-th position.*

**Definition 6.5** (Linear weights). *Let $\theta \in \mathcal{H}_\sigma$ and $k \in \{1, .., p\}$. We define the $k$-th feature weight of $\theta$, termed $\mathcal{L}_k$, as the projection on $G_{\delta_k}$:*

$$\mathcal{L}_k(\theta) \;=\; \langle \theta, G_{\delta_k} \rangle_{\mathcal{H}_\sigma}. \tag{42}$$

This quantity intuitively corresponds to the weight of a single-feature (the $k$-th) modulated by the squared exponential term.

**Definition 6.6** (Interaction weight). *Let $\theta \in \mathcal{H}_\sigma$ and let $k, l \in \{1, .., p\}$. We define the $k, l$-interaction weight of $\theta$, termed $\mathcal{I}_{k,l}$, as the projection on $G_{\delta_k+\delta_l}$:*

$$\mathcal{I}_{k,l}(\theta) = \langle \theta, G_{\delta_k+\delta_l} \rangle_{\mathcal{H}_\sigma}. \tag{43}$$

This quantity intuitively corresponds to the weight of an interaction term, i.e. the product of two genes, modulated by the squared exponential term. We can observe that, when $k = l$, the interaction terms corresponds to a quadratic term.

**Definition 6.7** ($I$-higher order weight). *Let $\theta \in \mathcal{H}_\sigma$ be an element of the Gaussian kernel RKHS. Let $I \in \mathbb{N}^p$ with $|I| = \sum_{i=1}^p I_i \geq 3$. The $I$-higher-order-interaction weight of $\theta$, termed $\mathcal{I}_I^+$, is defined as the projection on $G_I$:*

$$\mathcal{I}_I^+(\theta) = \langle \theta, G_I \rangle_{\mathcal{H}_\sigma}. \tag{44}$$

Using these three kinds of contributions, we can define the global contribution of linear, interaction and higher order interactions as follows.

**Definition 6.8** (Linear, interactions and higher order interactions contributions). *Let $\theta \in \mathcal{H}_\sigma$. We define the linear, interactions and higher-order interactions contributions, called $\mathcal{L}$, $\mathcal{I}$ and $\mathcal{I}^+$ respectively, as the squared norm of the associated vector weights, i.e.:*

$$\mathcal{L}(\theta) = \sum_{k=1}^p \mathcal{L}_k(\theta)^2, \quad \mathcal{I}(\theta) = \sum_{1\leq k\leq l\leq p} \mathcal{I}_{k,l}(\theta)^2 \quad \text{and} \quad \mathcal{I}^+(\theta) = \sum_{\substack{I\in\mathbb{N}^p \\ |I|\geq 3}} \mathcal{I}_I^+(\theta)^2. \tag{45}$$

#### 6.3 Computation of gene-level contribution (linear)

We now present how to compute the linear weights. These rely on the artificial samples (Definition 4.5) used for approximating the latent factor (Definition 4.6).

**Definition 6.9** (Offset matrices). *We define the two offset matrices $\mathcal{O}^X \in \mathbb{R}^M$ and $\mathcal{O}^Y \in \mathbb{R}^M$ as*

$$\mathcal{O}^X = \operatorname{diag}\left[\exp\left(-\frac{\|\widehat{x}_i^X\|^2}{2\sigma^2}\right)\right] \quad \text{and} \quad \mathcal{O}^Y = \operatorname{diag}\left[\exp\left(-\frac{\|\widehat{x}_i^Y\|^2}{2\sigma^2}\right)\right] \tag{46}$$

**Definition 6.10** (Model points matrices). *We define the two artificial sample matrices $\mathcal{A}^X \in \mathbb{R}^{M\times p}$ and $\mathcal{A}^Y \in \mathbb{R}^{M\times p}$ as:*

$$\mathcal{A}^X = \left[\widehat{x}_1^X, \, .. \, , \widehat{x}_M^X\right]^T \quad \text{and} \quad \mathcal{A}^Y = \left[\widehat{x}_1^Y, \, .. \, , \widehat{x}_M^Y\right]^T \tag{47}$$

**Theorem 6.11** (Computing individual contributions). *We have the following equalities, for all $t \in \{X, Y\}$:*

$$\begin{aligned}
\mathcal{L}_{fact}^t &= \left(\mathcal{L}_j\left(\theta_i^t\right)\right)_{\substack{1\leq i\leq d_t \\ 1\leq j\leq p}} &= \frac{1}{\sigma}\alpha^t \mathcal{O}^t \mathcal{A}^t. \\
\mathcal{L}_{SPV}^X &= \left(\mathcal{L}_j\left(s_i\right)\right)_{\substack{1\leq i\leq d_X \\ 1\leq j\leq p}} &= \frac{1}{\sigma}\gamma^X \mathcal{O}^X \mathcal{A}^X. \\
\mathcal{L}_{SPV}^Y &= \left(\mathcal{L}_j\left(t_i\right)\right)_{\substack{1\leq i\leq d_Y \\ 1\leq j\leq p}} &= \frac{1}{\sigma}\gamma^Y \mathcal{O}^Y \mathcal{A}^Y.
\end{aligned} \tag{48}$$

*Proof.* We first recall the second reproducing property of the kernel $K_\sigma$:

$$\forall f \in \mathcal{H}, \forall x \in \mathbb{R}^p, \quad \left\langle K_\sigma^G (x, \cdot), f \right\rangle_{\mathcal{H}_\sigma} = f(x). \tag{49}$$

Let $1 \le i \le d_t$ and $1 \le j \le p$. Using Definitions 4.6 Definition 6.5 and the aforementionned second reproducing property, we have:

$$\mathcal{L}_j\left(\theta_i^t\right) = \sum_{k=1}^{M} \alpha_{i,k}^t \left\langle K_\sigma^G\left(\widehat{x}_k^t, \cdot\right), G_{\delta_j}\right\rangle_{\mathcal{H}_\sigma} = \sum_{k=1}^{M} \alpha_{i,k}^t G_{\delta_j}\left(\widehat{x}_k^t\right). \tag{50}$$

Using the definition of the Gaussian basis functions (Definition 6.2), we obtain:

$$\mathcal{L}_j\left(\theta_i^t\right) = \sum_{k=1}^{M} \alpha_{i,k}^t \frac{\widehat{x}_{k,j}^t}{\sigma} \exp\left(-\frac{\left\|\widehat{x}_k^t\right\|^2}{2\sigma^2}\right), \tag{51}$$

which, put in matrix format, gives the desired result.
Same idea gives the other two results, using the expansion of Theorem 5.2 instead of Definition 4.6. ■

It follows from Theorem 6.11 that the global linear contribution can be computed as follows.

**Proposition 6.12** (Computing global contributions)**.** *We have the following equalities:*

$$\forall t \in \{X, Y\}, \forall i \in \{1, .., d_t\}, \quad \mathcal{L}\left(\theta_i^t\right) = \frac{1}{\sigma^2}\left(\alpha^t \mathcal{O}^t \mathcal{A}^t {\mathcal{A}^t}^T \mathcal{O}^t {\alpha^t}^T\right)_{i,i}. \tag{52}$$

$$\forall i \in \{1, .., d_X\}, \quad \mathcal{L}(s_i) = \frac{1}{\sigma^2}\left(\gamma^X \mathcal{O}^t \mathcal{A}^t {\mathcal{A}^t}^T \mathcal{O}^t {\gamma^X}^T\right)_{i,i}. \tag{53}$$

$$\forall i \in \{1, .., d_Y\}, \quad \mathcal{L}(t_i) = \frac{1}{\sigma^2}\left(\gamma^Y \mathcal{O}^t \mathcal{A}^t {\mathcal{A}^t}^T \mathcal{O}^t {\gamma^Y}^T\right)_{i,i}. \tag{54}$$

*Proof.* Immediate by combining Theorem 6.11 with Definition 6.8. ■

#### 6.4 Computation of interaction weights

**Definition 6.13** (Gene expression product matrix)**.** *We define the matrices $\overline{\mathcal{A}^X}$ and $\overline{\mathcal{A}^Y}$ as*

$$\begin{aligned} \overline{\mathcal{A}}^X &= \left[\mathcal{A}_{:,i}^X \circ \mathcal{A}_{:,j}^X\right]_{1 \le i \le j \le p} \in \mathbb{R}^{M \times \frac{p(p+1)}{2}} \\ \overline{\mathcal{A}}^Y &= \left[\mathcal{A}_{:,i}^Y \circ \mathcal{A}_{:,j}^Y\right]_{1 \le i \le j \le p} \in \mathbb{R}^{M \times \frac{p(p+1)}{2}}, \end{aligned} \tag{55}$$

*where $\circ$ is the Hadamard (piece-wise) product between two vectors.*

The product matrices presented in Definition 6.13 correspond to the products of columns of $\mathcal{A}^X$ and $\mathcal{A}^Y$. The indices $i$ and $j$ are ordered by first setting $i$ and then varying $j$ in increasing order, i.e. $(1,1), (1,2), .., (1,p), (2,2), .., (2,p), .., (p-1,p-1), (p-1,p), (p,p)$.
In order to keep track of the different interaction terms and avoid computing the same interaction terms twice, we introduce the following interaction indexing.

**Definition 6.14** (Interaction indexing)**.** *We define the interaction indexing function $\Upsilon$ as:*

$$\forall i, j \in \{1, .., p\}, \quad \Upsilon(i,j) = \begin{cases} (i-1)\left(p+1-\frac{i}{2}\right)+j & \text{if } i \le j \\ \Upsilon(j,i) & \text{otherwise} \end{cases}. \tag{56}$$

**Proposition 6.15.** *The interaction weights can be computed as follows, for $t \in \{X, Y\}$ and $k \in \{1, .., d_t\}$:*

$$\forall 1 \leq i < j \leq p, \quad \begin{aligned} \mathcal{I}_{i,j}\left(\theta_k^t\right) &= \frac{1}{\sigma^2}\left(\alpha^t \mathcal{O}^t \overline{\mathcal{A}}^t\right)_{k,\Upsilon(i,j)} \\ \mathcal{I}_{i,j}\left(s_k\right) &= \frac{1}{\sigma^2}\left(\gamma^X \mathcal{O}^X \overline{\mathcal{A}}^X\right)_{k,\Upsilon(i,j)} \\ \mathcal{I}_{i,j}\left(t_k\right) &= \frac{1}{\sigma^2}\left(\gamma^Y \mathcal{O}^Y \overline{\mathcal{A}}^Y\right)_{k,\Upsilon(i,j)} \end{aligned}. \tag{57}$$

#### 6.5 Interpretration in the Laplacian kernel

The orthonormal basis of the the Gaussian kernel of length-scale $\sigma$ relies on the Hilbertian structure of the Gaussian RKHS $\sigma$. However, the Gaussian kernel $K_\sigma$ and the Laplacian kernel $K_\sigma^L$ correspond to the two extreme of the Matérn family: with $\nu$ going to infinity for the Gaussian and to $\frac{1}{2}$ for the Laplacian kernel. We use this identity to transfer the feature weights from the Gaussian setting to the Laplacian setting, which we present here. This identification relies on two lemmas.

**Lemma 6.16.** *Let $k, l \in \{1, .., p\}$ with $k \neq l$, then*

$$\langle G_{\delta_k}, G_{\delta_l} \rangle_{\mathcal{H}_\sigma^L} = 0 \tag{58}$$

*Proof.* Let's denote by $\mathcal{F}[f]$ the Fourier transform of a function $f$. Following [Kimeldorf et al] [9] (Lemma 3.1), the inner-product of the Laplacian RKHS is computed as follows, for $f$ and $g$ two real functions:

$$\langle f, g \rangle_{\sigma^L} = \frac{1}{(2\pi)^{p/2} C_{\sigma,p}} \int_{\mathbb{R}^p} \mathcal{F}[f] \overline{\mathcal{F}[g]} \left(\frac{2}{\sigma^2} + 4\pi^2 \|\omega\|^2\right)^{\frac{1+p}{2}} d\omega, \tag{59}$$

with $\bar{\cdot}$ indicating complex conjugate. Noting that $\mathcal{F}[G_{\delta_k}](\omega) = -i\frac{\omega_k}{2\sigma^2}\exp\left(-\frac{\|\omega\|^2}{2\sigma^2}\right)$ by partial derivation, we can write:

$$\langle G_{\delta_k}, G_{\delta_l} \rangle_{\mathcal{H}_\sigma^L} = \frac{-1}{(2\pi)^{p/2} C_{\sigma,p}} \int_{\mathbb{R}^p} \omega_k \omega_l \left(\frac{2}{\sigma^2} + 4\pi^2 \|\omega\|^2\right)^{\frac{1+p}{2}} \exp\left(-\frac{\|\omega\|^2}{\sigma^2}\right). \tag{60}$$

The integrated function is odd with regard to the plane $\omega_k = 0$, so $\langle G_{\delta_k}, G_{\delta_l} \rangle_{\mathcal{H}_\sigma^L} = 0$. ■

**Lemma 6.17.** *Let $k_1, k_2, l_1, l_2 \in \{1, .., p\}$ with $(k_1, l_1) \neq (k_2, l_2), k_1 < l_1, k_2 < k_2$, then*

$$\left\langle G_{\delta_{k_1}+\delta_{l_1}}, G_{\delta_{k_1}+\delta_{l_1}} \right\rangle_{\mathcal{H}_\sigma^L} = 0 \tag{61}$$

*Proof.* If $k_1 \neq l_1$, we have:

$$\mathcal{F}\left[G_{\delta_{k_1}+\delta_{l_1}}\right] = -\frac{\omega_{k_1}\omega_{l_1}}{4\sigma^2} \exp\left(-\frac{\|\omega\|^2}{2\sigma^2}\right). \tag{62}$$

■

These two lemmas shows that the linear and interaction features – except quadratic terms – form an orthogonal family of the Laplacian kernel RKHS. Their norm, however, is not unit and computation thereof is analytically challenging and computationally untractable due to the high dimensionality of the integration problem. In order to correct and obtain a unit orthogonal family, we correct by using the linear and interaction contributions from the Gaussian kernel (Definition 6.8). Specifically, for a function $f$, we multiply all the linear terms by $\frac{\mathcal{L}(f)}{\mathcal{L}^L(f)}$ where $L$ superscripts refers to the contribution in the Laplacian kernel (no super-script for the gaussian kernel). We similarly multiply all interaction terms by $\frac{\mathcal{I}(f)}{\mathcal{I}^L(f)}$.

### 7 Algorithm

**Algorithm 1** Sobolev Alignment

**Require:** Datasets $\mathcal{X}$ and $\mathcal{Y}$, scVI parameters, number anchors $M$, number of artificial points $N$, penalizations $\lambda_X$ and $\lambda_Y$, Matérn kernel $K_{\nu,\sigma}$.

Train scVI (VAE) model $\mathcal{M}_X$ on $\mathcal{X}$ ($d_X$ hidden neurons).
Train scVI (VAE) model $\mathcal{M}_X$ on $\mathcal{Y}$ ($d_Y$ hidden neurons).
$Z_X \leftarrow N$ vectors sampled from $\mathcal{N}(0, I_{d_X})$.
$\widehat{X}_X \leftarrow$ decoding of $Z_X$ using decoder of $\mathcal{M}_X$
$\widehat{Z}_X \leftarrow$ encoding of $\widehat{X}_X$ using encoder of $\mathcal{M}_X$
$Z_Y \leftarrow N$ vectors sampled from $\mathcal{N}(0, I_{d_Y})$.
$\widehat{X}_Y \leftarrow$ decoding of $Z_Y$ using decoder of $\mathcal{M}_Y$.
$\widehat{Z}_Y \leftarrow$ encoding of $\widehat{X}_Y$ using encoder of $\mathcal{M}_Y$.
$\theta_1^X, ...\theta_{d_X}^X \leftarrow$ KRR models between $\widehat{X}_X$ (input) and $\widehat{Z}_X$ (label).
$\theta_1^Y, ...\theta_{d_Y}^Y \leftarrow$ KRR models between $\widehat{X}_Y$ (input) and $\widehat{Z}_Y$ (label).
$\alpha^X \leftarrow$ Sample coefficients of $\theta_1^X, ...\theta_{d_X}^X$.
$\alpha^Y \leftarrow$ Sample coefficients of $\theta_1^Y, ...\theta_{d_Y}^Y$.
$K_X, K_Y, K_{XY} \leftarrow$ Kernel matrices using $K_{\nu,\sigma}$ on $\widehat{X}_X$ and $\widehat{X}_Y$. ▷ Definition 4.8.
$\widetilde{\mathbf{M}}_X \leftarrow \alpha^X K_X {\alpha^X}^T$.
$\widetilde{\mathbf{M}}_Y \leftarrow \alpha^Y K_Y {\alpha^X}^Y$.
$\widetilde{\mathbf{M}}_{X,Y} \leftarrow \alpha^X K_{X,Y} {\alpha^Y}^T$.
$\mathbf{M} \leftarrow \widetilde{\mathbf{M}}_X^{-1/2} \mathbf{M}_{X,Y} \widetilde{\mathbf{M}}_Y^{-1/2}$. ▷ Definition 4.10.
$U, \Sigma, V \leftarrow$ SVD decomposition of $\mathbf{M}$ ▷ $\mathbf{M} = U\Sigma V^T$.
$\gamma^X \leftarrow U^T \widetilde{\mathbf{M}}_X^{-1/2} \alpha^X$.
$\gamma^Y \leftarrow V^T \widetilde{\mathbf{M}}_Y^{-1/2} \alpha^Y$. ▷ Theorem 5.2.

### 8 Glossary

| Symbol | Meaning | Reference |
|---|---|---|
| $p$ | Number of genes (features). | |
| $n_X$ and $n_Y$ | Number of source (cell-lines) and target (tumors) samples. | |
| $\mathbb{R}$ | Real numbers. | |
| $I_d$ | Identity matrix of size $d$. | |
| diag | Diagonal matrix. | |
| $\langle \cdot, \cdot \rangle$ | Inner-product. | |
| $\mathcal{N}(\mu, \Sigma)$ | Multivariate normal distributio, $\mu$: mean, $\Sigma$: covariance. | |
| $\mathcal{X}_X$ and $\mathcal{X}_Y$ | Source and target datasets, of sizes $n_X \times p$ and $n_Y \times p$. | |
| $K_\sigma^L$ | Laplacian kernel, $\sigma$:lengthscale. | Definition 2.4 |
| $K_{\nu,\sigma}^M$ | Matérn kernel, $\nu$:smoothness, $\sigma$:lengthscale. | Definition 2.5 |
| $K_\sigma^G$ | Gaussian kernel, $\sigma$:lengthscale. | Definition 2.6 |
| $\Gamma$ | Gamma function | |
| $K_\alpha$ | Modified Bessel function of second kind of order $\alpha$. | |
| $L_2(\mathbb{R}^p)$ | Space of continuous integrable functions. | Definition 2.9 |
| $W_2^s(\mathbb{R}^p)$ | Sobolev space of order $s$. | Definition 2.10 |
| $\mathcal{H}_{\nu,\sigma}$ | RKHS associated to the Matérn kernel $K_{\nu,\sigma}^M$. | Proposition 2.12 |
| $d_X$ and $d_Y$ | Number of source and target latent variables. | Definition 3.2 |
| $\mu_1^X, .., \mu_{d_X}^X$ | Mean embedding function for source VAE. | Definition 3.2 |
| $\Sigma_1^X, .., \Sigma_{d_X}^X$ | Standard-deviation embedding function for source VAE. | Definition 3.2 |
| $f_1^X, .., f_{d_X}^X$ | Encoding (mean) functions for cell lines scVI model. | Definition 4.1 |
| $z_1^X, .., z_N^X$ | Points randomly sampled from noise (cell). | Definition 4.5 |
| $\widehat{x}_1^X, .., \widehat{x}_N^X$ | Decoded values of $z_1^X, .., z_N^X$ using cell line scVI model | Definition 4.5 |
| $\widehat{z}_1^X, .., \widehat{z}_N^X$ | Encoded values of $\widehat{x}_1, .., \widehat{x}_N^X$ using cell line scVI model | Definition 4.5 |
| $\widehat{\theta}_k^t$ | KRR approximations for encoding functions $\widehat{f}_k^t$ | Definition 4.6 |
| $M$ | Number of anchor points (Falkon approximation) | Proposition 4.4 |
| $\alpha^X$ and $\alpha^Y$ | Sample weights for KRR approximations | Definition 4.6 |
| $\widetilde{\mathbf{M}}_{XY}$ | Un-normalized cosine similarity matrix | Definition 4.10 |
| $\widetilde{\mathbf{M}}_X$ and $\widetilde{\mathbf{M}}_Y$ | Inner-product matrices between latent factors | Definition 4.10 |
| $K_X, K_Y$, and $K_{XY}$ | Similarity matrices (between samples) | Definition 4.8 |
| $\mathbf{M}$ | Cosine similarity matrix | Definition 4.10 |
| $\widehat{d}$ | Number of principal vectors (PVs) | Definition 5.1 |
| $s_1, ..., s_{\widehat{d}}$ | Source principal vectors | Definition 5.1 |
| $t_1, ..., t_{\widehat{d}}$ | Target principal vectors | Definition 5.1 |
| $\gamma^X, \gamma^Y$ | Sample-weights for source and target PVs | Theorem 5.2 |
| $G_I, I \in \mathbb{N}^p$ | Gaussian basis function | Definition 6.2 |
| $\mathcal{L}_k, k \in \{1, .., p\}$ | Linear weights | Definition 6.5 |
| $\mathcal{I}_{k,l}, k, l \in \{1, .., p\}$ | Interaction weights | Definition 6.6 |
| $\mathcal{O}^X, \mathcal{O}^Y$ | Offset matrices | Definition 6.9 |
| $\mathcal{A}^X, \mathcal{A}^Y$ | Model points matrices | Definition 6.10 |
| $\overline{\mathcal{A}}^X, \overline{\mathcal{A}}^Y$ | Gene expression products matrix | Definition 6.14 |
| $\Upsilon$ | Interaction indexing | Definition 6.13 |
| $\mathcal{F}$ | Fourier transform. | |

### References

[1] Jean-Philippe Vert, Koji Tsuda, and Bernhard Schölkopf. *Kernel methods in computational biology.* MIT Press, 2004.

[2] Bernhard Schölkopf and Alexander J. Smola. *Learning with kernels.* 2002.

[3] John Shawe-Taylor and Nello Cristianini. *Kernel Methods for Pattern Analysis.* 2004.

[4] Motonobu Kanagawa, Philipp Hennig, Dino Sejdinovic, and Bharath K Sriperumbudur. Gaussian Processes and Kernel Methods: A Review on Connections and Equivalences. pages 1–64, 2018.

[5] Bertil Matern. *Spatial Variation.*, volume 36. 1970.

[6] Diederik P. Kingma and Max Welling. Auto-encoding variational bayes. *2nd International Conference on Learning Representations, ICLR 2014 - Conference Track Proceedings*, (Ml):1–14, 2014.

[7] Romain Lopez, Jeffrey Regier, Michael B. Cole, Michael I. Jordan, and Nir Yosef. Deep generative modeling for single-cell transcriptomics. *Nature Methods*, 15(12):1053–1058, 2018.

[8] Ingo Steinwart, Don Hush, and Clint Scovel. An Explicit Description of the Reproducing Kernel Hilbert Spaces of Gaussian RBF Kernels. *IEEE Transactions on Information Theory*, 52(10):4635–4643, 2006.

[9] George S. Kimeldorf and Grace Wahba. A Correspondence Between Bayesian Estimation on Stochastic Processes and Smoothing by Splines. *The Annals of Mathematical Statistics*, 41(2):495–502, 1970.
